# Extracellular matrix stiffness modulates nuclear lamina organisation and sets nuclear conditions for PRC2 repression

**DOI:** 10.1101/2025.09.07.674693

**Authors:** Alexia Pigeot, Martin Rey-Millet, Amal Zine El Aabidine, Antonio Trullo, Lea Costes, Manoel Manghi, Kerstin Bystricky, Jean-Christophe Andrau, Cyril Esnault

**Affiliations:** Institut de Génétique Moléculaire de Montpellier, University of Montpellier, CNRS-UMR 5535, 1919 Route de Mende, 34293 cedex 5, Montpellier, France; Université de Toulouse, CNRS, Centre de Biologie Integrative, CBI, MCD, Toulouse, France; Université de Toulouse, CNRS, Laboratoire de Physique Théorique, IRSAMC, Toulouse, France

## Abstract

Capability of cells to respond to tissue-level elasticity has important physiological and pathological implications. Stiffening of the extracellular matrix (ECM) promotes invasive behaviour of cancer cells, supports the transformation of fibroblasts into cancer-associated fibroblasts and primes stem cell differentiation programs. Here, we investigated how ECM stiffness modulates the Nuclear Lamina (NL) and its impact on gene expression programs, epigenetic marking and 3D genome organisation. By combining hydrogel cell culturing of primary fibroblasts, genomics and super-resolution microscopy, we found that ECM stiffness modifies composition of the NL, modulates long range chromatin interactions, induces changes in chromatin motion and regulates thousands of genes. We identified a specific set of genes coding proteins involved in pathways related to mechanical adaptation such as adhesion and signalling. These genes harbour an apparent bivalent chromatin signature and are expressed under soft condition while repressed in stiff condition through Polycomb Repressive Complex 2 (PRC2). We found that this stiffness-specific repression is tempered by mechano-transduction and the NL. This work uncovers mechano-dependent NL composition, changes in 3D genome organisation and in chromatin motion which underlie adaptative gene expression programs controlled through PRC2.

## Introduction

Mechano-sensation controls morphogenesis, tissue maintenance and regulates cellular homeostasis, behaviour and commitments. Mechanical cues arise from both active forces such as shear, compression or stretch and from forces generated by the cell itself resisting to viscoelastic properties of the extracellular matrix (ECM) or the surrounding cells ^1,2^. Modifications of the ECM stiffness is of physiological and pathological importance. It regulates mesenchymal and epidermal stem cell fates ^3,4^, invasiveness of cancer cells ^5^ and influences fibroblast transformation into cancer associated fibroblasts (CAFs) ^6,7^. Mechanical properties of the ECM can fluctuate from tissues mechanics ^8^, during tumour growth ^5^ or with aging ^9^. Generally, long-term cell function is sensitive to environmental cues which remodel the epigenetic status and expression programs ^10^. But, how are epigenetics and gene expression regulation established in response to mechanical cues remain largely elusive: in particular how mechanical transmission of forces to the Nuclear Lamina (NL) impacts chromatin properties and genome organisation, as well as the physiological relevance of such modifications and the definition of gene sets sensitive to nuclear mechanics.

Cell adhesion to ECM and mechano-transduction involve focal adhesion (FA) complexes and actin cytoskeleton reorganisation ^11^. Actin polymerisation in response to increasing stiffness of the ECM alters the rates of globular actin (G-actin) and filamentous actin (F-actin) pools. A decrease in the limited G-actin pool leads to Myocardin-Related Transcription Factors (MRTFs) activation which in turn co-activate the Serum Response Factor (SRF) and its expression programs ^12–15^. Similarly, increased actomyosin cytoskeleton tension promotes YAP/TAZ transcription factor activity ^16^. In addition to these well-established transcriptional systems, a direct implication of the NL in gene expression program regulation in response to mechanical cues remains largely unknown.

The cytoskeleton is physically connected to the NL through the Linker of Nucleoskeleton and Cytoskeleton (LINC) complex, composed of SUN and Nesprin proteins ^17^, which transmit mechanical forces to the nucleus ^18,19^. The NL is a meshwork of nuclear intermediate filaments lining the inner part of the nucleus consisting of Lamin A and C isoforms, encoded by *LMNA* gene, and Lamin-B1 and B2 respectively encoded by *LMNB1 and LMNB2*. The latter are ubiquitously expressed while Lamin-A/C expression is acquired through cell differentiation and development ^20,21^. Lamin-A/C expression tunes with ECM stiffness ^8^ and acts as a nucleoskeleton by regulating nuclear mechanics ^22^.

Mechanical cues may also directly act at the chromatin level. In the nucleus, the genome is organized in functional domains exhibiting features that show preferential chromatin interactions delimiting Topologically Associating Domains (TADs) ^23–26^. Large chromatin regions are also found at the nuclear periphery in association with the NL creating the Lamina Associating Domains (LADs). LADs mainly contain transcriptionally inactive genes and are enriched in repressive histone marks such as H3K27me3 and H3K9me2/3 ^27–31^. Some LADs are constitutively found and shared between various cell types (cLAD) while facultative LADs (fLADs) are cell-type specific and acquired during cell differentiation ^32,33^. Genome organisation is not static: local chromatin fibre folding varies from cell to cell and protein complexes stochastically coalesce and diffuse. Chemical stimuli or external forces could modulate epigenetic states. Mechanical transmission of forces to the NL and to chromatin could thus impact chromatin dynamics. ECM elasticity tunes the expression of NL components such as Lamin-A types or the Lamin-B Receptor (LBR) inclusion and affects NL properties ^8,34^. Hence, mechano-sensation and nuclear mechanics may have a direct impact on gene expression through the NL.

Finally, mechanical cues also signal on epigenetic marking and histone modifications. Histone modifications characterise promoters and regulatory elements in active, inactive or poised states ^35^. H3K27me3 is a mark deposited by the Polycomb Repressive Complex 2 (PRC2) and associated with gene repression ^36^. Acute mechanical strain leads to global gene repression with increased H3K27me3 marking, in an actin dependent manner ^37,38^. H3K4me3 is found at most promoters which globally correlates with the level of transcription but which can also be found at inactive loci ^39,40^. Both marks can be associated at the same genomic locations, defining bivalent promoters that are generally poised for transcription ^41,42^. Bivalent promoters were first described in mouse embryonic stem (ES) cells and found to mark developmental genes ^43–47^. They are not restricted to pluripotent cells and were observed in differentiated cells where their roles remain uncharacterised ^45–48^. Interestingly, recent works associate ECM niche stiffening with reduced ability to activate bivalent genes for efficient self-renewal and differentiation of hair follicle stem cells during aging ^9^.

To uncover the impact of ECM stiffness on chromatin organisation and gene expression, we cultured primary human fibroblasts (BJ) on polyacrylamide hydrogels mimicking physiological stiffness: 0.5kPa (soft), observed near the vascular parts of the bone marrow or of soft primary solid tumours and 50kPa (stiff), found at the proximity of bones or in fibrotic tumours ^49–51^. We found that nuclear flattening and stiffening following exposure to stiff substrate reduces long-range 3D chromatin contacts and chromatin mobility especially in areas close to the nuclear periphery. In addition, we identified 310 genes with apparent bivalent chromatin more frequently associated to the NL and whose repression under stiff condition requires NL components. Our study suggests that nuclear reorganisations driven by mechanical cues regulate gene expression not only through chromatin landscape modifications but also through the regulation of chromatin motion.

## Results

### Extracellular matrix stiffness tunes nuclear lamina organisation

Hydrogels have emerged in the last decades for cell culture, they mimic salient elements of native extracellular matrices (ECMs), have tuneable mechanics ranging from soft to stiff tissues, and can support cell adhesion and protein sequestration ^52^. We have used hydrogels of 0.5kPa (soft) and 50kPa (stiff) to study responses to extracellular matrix stiffness in primary fibroblast (BJ cells) and to mimic mechanical conditions to which fibroblast cells are exposed to (Fig. 1a). Using confocal microscopy, we confirm cytoskeleton modifications with increased cell surface, enhanced staining of actin stress fibres (phalloidin) and increased adhesion foci marked with Vinculin under stiff hydrogels (Fig. 1b).

**Figure 1:**
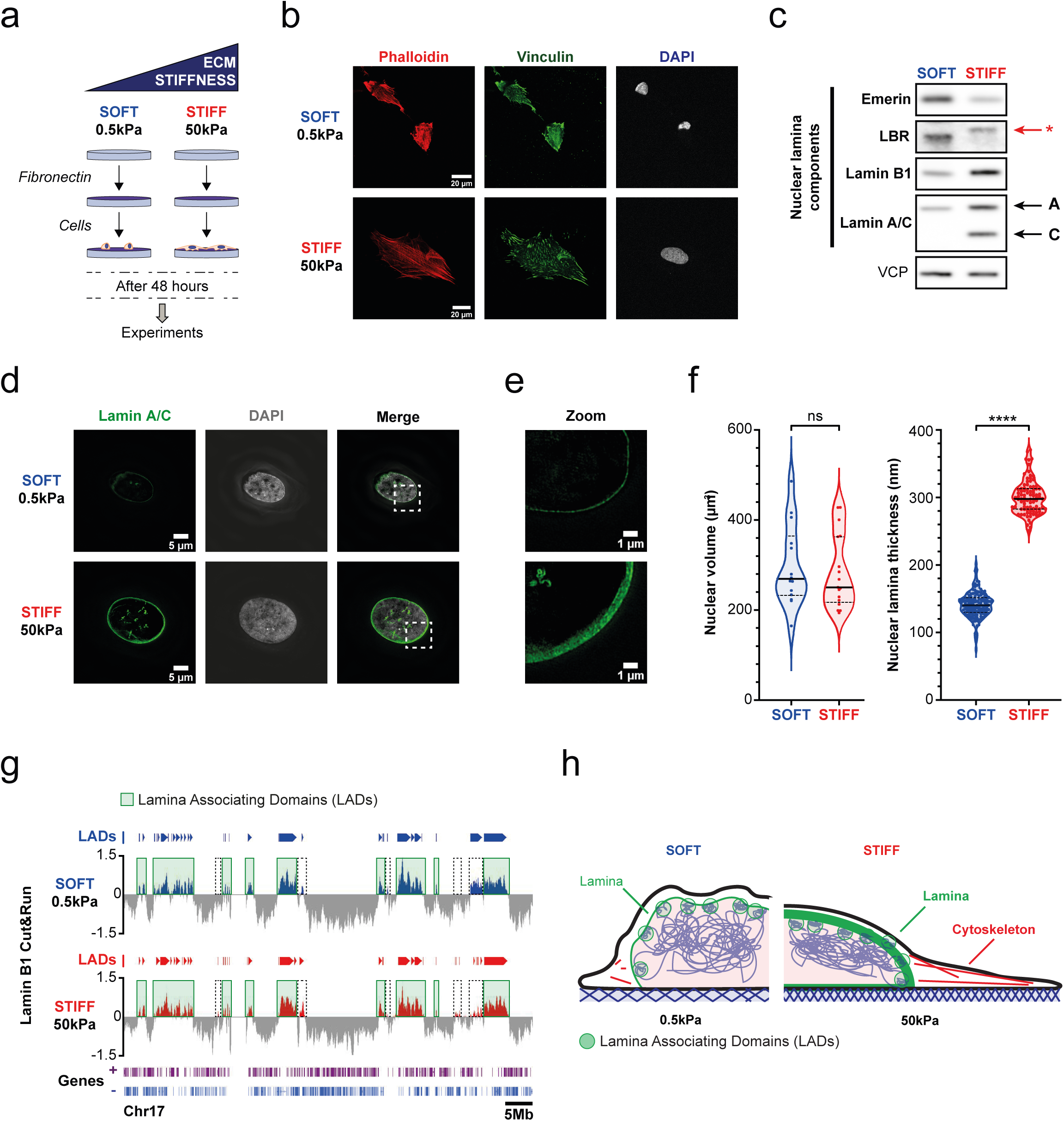
ECM stiffness regulates cell morphology and nuclear lamina organisation without major alteration of lamina associating domains distribution. a. Experimental setup. b. Representative confocal images of primary BJ fibroblasts plated on SOFT (top) or STIFF (bottom) hydrogels. Actin stress fibers are labelled with phalloidin (red), focal adhesions are visualised by vinculin staining (green) and nuclei are labelled with DAPI staining (white). Scaling bar: 20µm. c. Western blot of Nuclear Lamina (NL, Lamin A/C and Lamin B1) and NL associated components (LBR and Emerin) at different hydrogel stiffness in primary BJ fibroblasts. Lamin A and C isoforms are highlighted with black arrows. An asterix illustrates an observed shift in LBR size depending on the stiffness conditions. Loading control: VCP (Valosin-Containing Protein). See Figure S1a for quantifications. d. Representative super-resolution microscopy images of nuclei of primary BJ fibroblasts plated either on SOFT (top) or STIFF (bottom) hydrogels. NL component Lamin A/C are in green and nuclei are labelled with DAPI (white). Right: merge of Lamin A/C and DAPI channels. Scaling bar: 5µm. e. Zoom of Lamin A/C signal from dashed white scared region in e. Scaling bar: 1µm. f. Quantifications of nuclear volumes and NL thickness from super-resolution images. Blue: values for cells plated on SOFT, red: values for cells plated on STIFF. Statistical tests: Mann-Whitney. Median and quartiles are shown as dashed lines. g. Representative IGB screenshots of Lamin B1 Cut&Run signal normalised over IgG negative control for the whole chromosome 17 in SOFT (top) or STIFF (bottom) condition. Called lamina associating domains (LADs) are illustrated on the top of each track and highlighted with light green rectangles. Dashed black squares illustrate genomic region differentially associated to Lamin B1 in different stiffness conditions. h. Model illustrating observed cell and nuclear changes.

Nuclear shape is also sensitive to ECM stiffness as previously shown ^8^ with small and wrinkled nuclei on soft matrix, and smoothed-out and flattened nuclei on stiff matrix (Fig. 1b). We confirm and extend this observation using western blotting of the main NL components. We found increased levels of Lamins A/C and B1 under stiff condition while Emerin and LBR signals are stronger under soft condition (Fig. 1c, Extended Data Fig. 1a). We also note a change in gel mobility of LBR indicating further post translational modifications. Using super resolution microscopy, we then show that ECM stiffness promotes the NL thickening. The NL width increases from ∼140nm on soft hydrogels to ∼300nm on stiff hydrogels (Fig. 1d-f). ECM stiffness however does not affect nuclear volume of fibroblasts which are flattened under stiff condition (Fig. 1f).

To then investigate if the observed modifications of the NL organisation influence the lamina association with DNA, we scored genome wide LADs under soft and stiff conditions using Cut&Run, targeting Lamin B1 (Fig. 1g). Our results show that LADs size ranges between 40kb and 8Mb with a median size of about 250kb and are linked to genes with lower expression levels measured by total RNA-seq. These analyses are consistent with previous studies ^27^ (Extended Data Fig. 1b-c). We found that most LADs remain unchanged (∼97%) in response to matrix stiffness and only 236 domains display mechano-specific tethering (81 domains associate with the lamina under soft and 155 under stiff condition - Fig. 1g, Extended Data Fig. 1d-e, Extended Data Table 1). Our results demonstrate that the nuclear shape, the NL organisation, its properties and its composition are modulated in response to matrix stiffness with only minor changes in LAD identity (Fig. 1h).

### Extracellular matrix stiffness promotes changes of chromatin motion at the nuclear periphery and shapes 3D compartment contacts

Chromosomes are long fibers with structural dynamics that can be investigated from the angle of their polymer nature ^53^. Within the nucleus, the genome is 3D organised in space and in time, defining chromosome territories, compartments, TADs and loops, all being closely related to transcription program regulations. The NL also contributes to this organisation and influences gene expression through LADs ^54–56^. Thus, to investigate and quantify the impacts of ECM stiffness variations and of the modifications observed in the NL, in turn, influence chromatin motion, we used high-resolution diffusion mapping (HiD). HiD was developed to study motion of densely distributed fluorescent molecules such as chromatin labelled by incorporation of H2B-GFP ^57^. Time-lapse fluorescent microscopy is performed at 200 ms intervals over 1 - 3 minutes. HiD integrates dense optical flow methods to reconstruct trajectories of imaged fluorescent molecules at sub-pixel precision. In the subsequent steps, it fits the experimental Mean Square Displacement (MSD) of each trajectory with an anomalous diffusion model (MSD(t)=At^α with A=DT0^1-α), used to describe chromatin motion, allowing the extraction of biophysical parameters describing the dynamic behaviour of chromatin. In this way, diffusion constants (A) and anomalous exponents (α) are estimated for each pixel, allowing the generation of two-dimensional maps that depict chromatin dynamics at single-pixel resolution in living cells. (Fig. 2a). Anomalous exponents α inform on diffusion modes with values <1 indicating subdiffusion due to crowding or retention, equal to 1 suggesting Brownian motion, and >1 reflecting superdiffusion or directed motion which can result from active cellular processes (Fig. 2b). Here, we analysed local chromatin motion by HiD distinguishing two zones within nuclei of cells grown on soft vs stiff hydrogels. MSDs and anomalous exponents were determined within the most peripheral rim of 840 nm and compared to the ones assessed in the internal nuclear space (Fig. 2c). Importantly, HiD is not sensitive to the substrate as MSDs extracted from fixed cells on soft and stiff hydrogels are confounded. We found that chromatin motion differs between stiff and soft conditions and between the nuclear interior and the periphery. First, in the nuclear interior, chromatin motion is subdiffusive (α <1) and greater in cells grown under soft compared to those under stiff condition. However, at the nuclear periphery, the MSD curve under soft condition displays a distinct shape and rises above that of the curve determined under stiff condition, indicating enhanced chromatin motion likely driven by different underlying processes (Fig. 2c). Indeed, at the periphery, α>1 (1.1, SEM=0.03 ; active motion) in soft condition while α remains below 1 (subdiffusion) under stiff condition (Fig. 2d). This active motion at the nuclear periphery of soft matrix growth cells likely stems in part from movements of the nuclear envelope permitted by a less rigid NL. This motion is constrained following NL thickening in response to mechanical cues in stiff condition.

**Figure 2:**
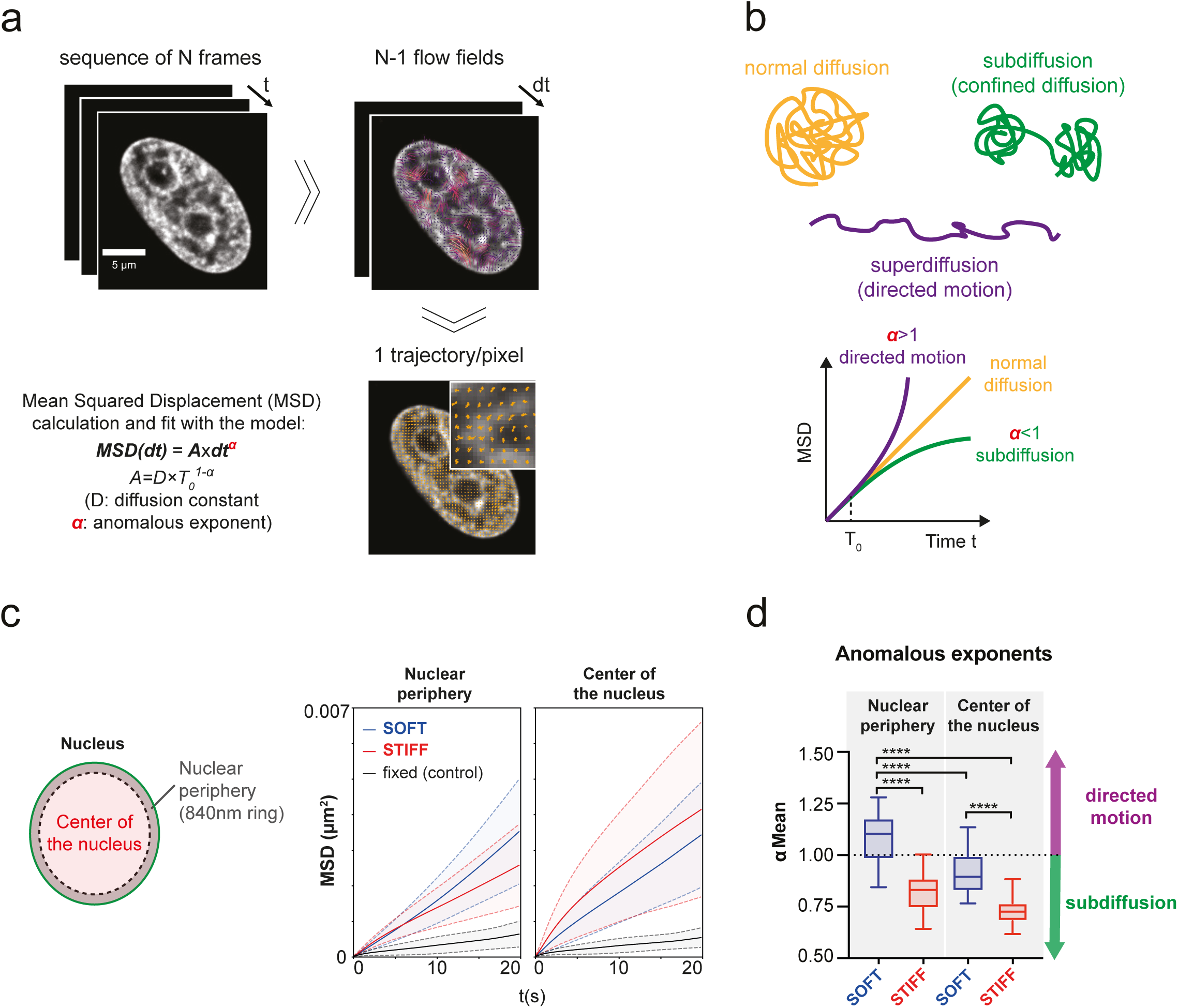
SOFT matrix promotes directed chromatin motion at the nuclear periphery. a. Hi-D analysis workflow. b. Scheme representing anomalous exponent values (α) inferred from HiD. c. Chromatin Mean Square Displacement (MSD) for chromatin located at nuclear periphery or in the center of the nucleus. Blue: MSD from cells plated on SOFT; red: MSD from cells plated on STIFF; black: MSD from fixed cells plated either on SOFT or STIFF. Dashed lines represent the standard deviations. d. Anomalous exponent values (α) depending on nuclear region (center or nuclear periphery or on stiffness. Statistical tests are one-way anova.

To then further infer how the NL modifications and changes in chromatin motion impact 3D genome architecture, we performed HiC experiments on cells grown on soft or stiff hydrogels (Fig.3a). Analysis of interaction frequencies indicates that long distance interactions are favoured under soft matrix growth condition in which the NL is thinner and chromatin motion is less constrained (Fig.3b, Wilcoxon test: 1.5e-148). We thus derived 3D interacting maps for each condition and differential contact frequencies *all to all* loci. The differential map confirmed the increase in long range contact frequencies on soft hydrogels (Fig.3c-d). Hence, we compared the 3D interaction maps with our LAD mapping and observed that structures that show differential interaction frequencies are largely defined by LAD positions (Fig.3c-d). To further validate and visualise these modifications in the 3D genome architecture, we used a 3D genome reconstruction contacts tool that recreates spatial distances and three-dimensional genome structures from observed contacts between genomic loci^58^. In this reconstruction, the distance between the dots represents the probability of interaction between the genomic locations: the closer they are the most frequently they are interacting. We show our results at chromosomes 11 to 20 to avoid overcrowded display and confirmed that cells growing under soft matrix have enhanced 3D interactions (Fig.3e). Our datasets support the notion that ECM stiffness impacts 3D genome organisation. Notably, on stiff matrix, the NL is thickening and chromatin motion in its vicinity is constrained. The changes in chromatin motion mode between the two conditions correlates with modifications in 3D genomic contacts largely defined by LADs. Overall, chromatin interactions seem favoured under soft growth “permissive” condition.

**Figure 3:**
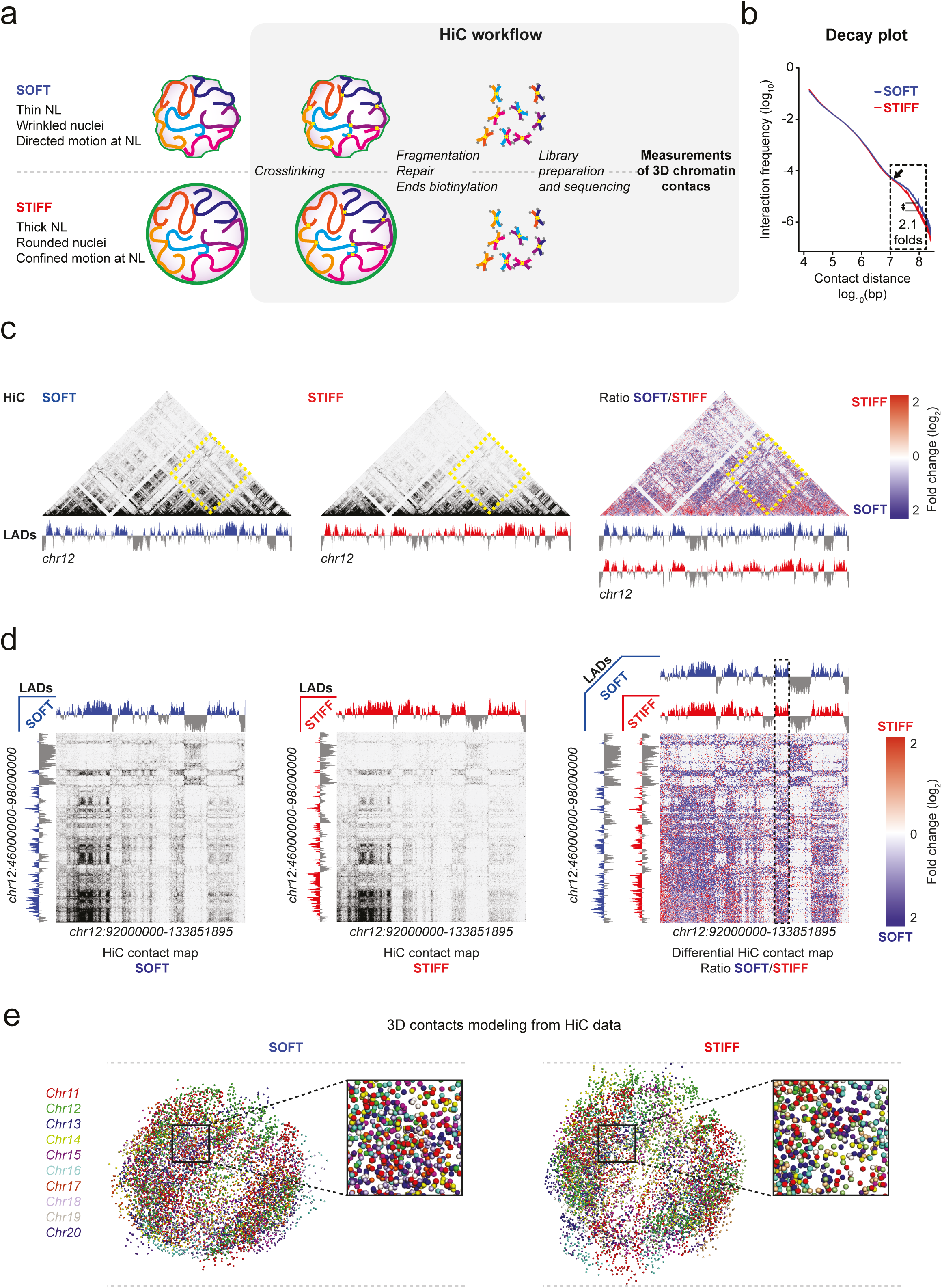
Long range 3D chromatin contacts are favoured under SOFT condition. a. Hi-C workflow. b. Decay plot representing interactions frequencies depending on contact distances. Dashed rectangle: highlight the increase of interaction frequencies in SOFT of long distance chromatin regions. c. Hi-C maps illustrating contact frequencies over the all chr12 and LADs signals from LaminB1 Cut&Run in cells plated on SOFT (left) and STIFF (middle) hydrogels. Right: differential Hi-C map. d. Zoom on the region defined by the dashed yellow rectangle in (c). e. 3D genome contact models based on Hi-C maps from each stiffness condition for chromosomes from 11 to 20.

### Extracellular matrix stiffness sets adaptative gene expression programs

To investigate the influence of the ECM stiffness on gene expression programs, we then performed total RNA sequencing on primary fibroblasts subjected to soft and stiff hydrogels. We found that gene expression is changing to large levels in response to these mechanical stimuli. Genes such as *MMP11* and *CYR61* display very specific expression patterns depending on which stiffness the cells are exposed to (Fig. 4a). In total over 30% of the human annotated genes we analysed show significant ECM stiffness-dependent expression levels (5973/19238 with 2804 and 3069 more expressed on soft or on stiff condition respectively, adjusted p-value < 0.01; Fig. 4b, Extended Data Table 2). We further confirmed this observation by specifically quantifying precursor RNA using intronic reads on 1698 genes with sufficient read coverage to be called using the DESEQ algorithm (adjusted p-value < 0.01 ; Extended Data Fig. 2a-c, Extended Data Table 2).

**Figure 4:**
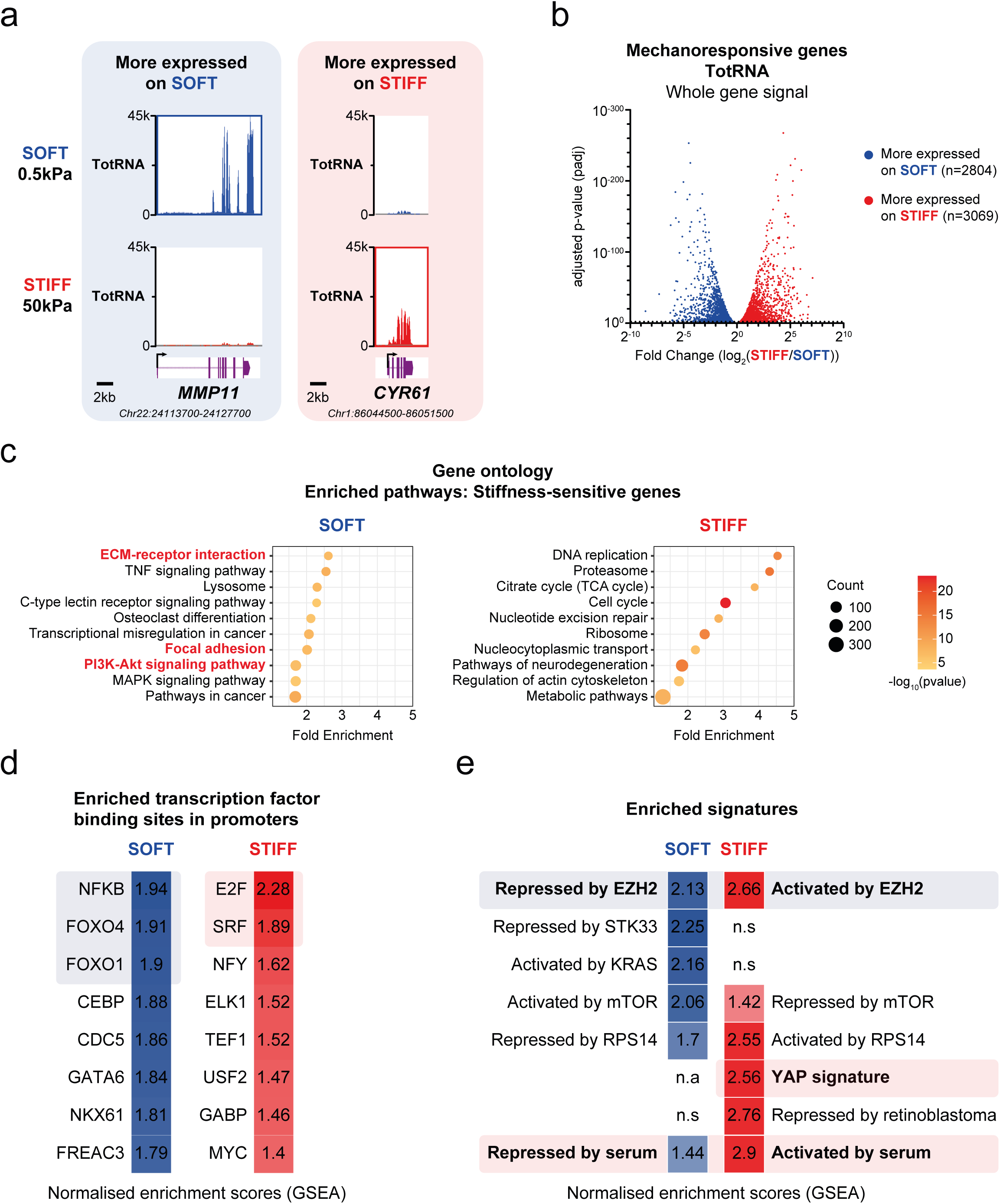
Substrate stiffness drives cell adaptation through modifications of gene expression programs. a. IGB screenshots of total RNA signal on stiffness specific mechano-responsive genes. Total RNA signal from cells plated on SOFT (top) and STIFF (bottom) hydrogels are in blue and red respectively. b. Volcano plot of differentially expressed genes at total RNA levels between stiffness conditions (DESeq package; threshold: padj < 0.01, Fold-Change: 1). Genes more expressed on SOFT are in blue and those more expressed on STIFF hydrogels are in red. c. Gene ontology (GO) analysis of differentially expressed genes using DAVID. d. Enriched transcription factor binding sites in promoters of differentially expressed genes identified (GSEA, TFT module). Blue: enriched in SOFT. Red: enriched in STIFF hydrogels. e. Identification of enriched gene signatures in cells plated on SOFT (blue) or STIFF (red) hydrogels using GSEA.

Then, we used gene ontology to identify the biological functions of genes with differential expression. Despite that gene sets displaying mechano-dependent expression patterns are very large, our gene ontology analyses were still able to highlight adaptative gene groups in our lists. Enriched groups in soft growth condition embrace regulatory transcription factors, signalling pathways, ECM regulators and components and negative regulator of cell proliferation, while on stiff, house-keeping genes are over-represented such as cell division, translation and DNA repair (Fig. 4c, Extended Data Fig. 2d). We then used the GSEA TFT module to explore transcription factors that are at play in mechano-transduction. This module catalogues motifs highly conserved in promoters of four mammalian species (human, mouse, rat and dog ; Fig. 4d) ^59^. We found SRF and E2F motifs as highly significant in our stiff gene signature which is consistent with SRF role as cytoskeleton and mechano-regulator ^2,13^ and with expression programs of cell division regulating genes. We also identified NFKB and Foxo family member enriched at promoters of the soft gene signature, pointing to potential input of unknown signalling programs in mechano-sensation. Finally, we used GSEA to interrogate enriched signatures across available expression datasets (Fig. 4e). Our main results comprise the YAP and, again, the serum response signatures which are known to involve cytoskeleton remodelling and to overlap with transduction of mechanical forces ^13,16^. Additionally, we found very high correlation scores between our signatures and PRC2-dependent genes in fibroblasts ^60^. In fact, sets of genes that are down regulated upon EZH2 knock-down, indicative of an activating role of EZH2 through possible indirect or non-canonical role of PRC2, are also enriched in our stiff signature. Furthermore, genes that are derepressed by inactivation of PRC2 components in fibroblasts ^60^ strongly overlap with the soft signature. These results strongly suggest that PRC2-dependent repression is impaired under soft growth condition and that EZH2, through an indirect or non-canonical role, supports the expression of some genes under stiff condition. Taken together, our transcriptome results indicate major reprogramming of gene expression during the transition from stiff to soft growth conditions associated with signalling and more wide-spread functions in soft and stiff media, respectively. Our data also suggest an involvement of PRC2 in the regulation of this transition.

### Soft matrix impedes PRC2-dependent repression of genes associated with H4K4me3 and H3K27me3 marking

Histone modifications regulate physical properties of chromatin and are important players in the regulation of gene expression ^61^. Chromatin accessibility remains an essential step for gene expression and promoter activity ^62^. Furthermore, our ontological analysis suggests a role of PRC2 in gene regulation in response to ECM stiffness. We thus used ATAC-seq to investigate how ECM stiffness and mechano-transduction shape the chromatin landscape and promoter opening of differently expressed genes between soft and stiff hydrogels (Fig. 5a). Our results show no major differences between the two conditions despite different gene activities. They are however consistent with known properties of mammalian promoters which display open chromatin structures independently of transcription ^63^. Histone modifications characteristic of active promoters or poised promoters such as H3K4me3 in association or not with H3K27me3 are proposed to maintain promoter in an open state ^42,64^. Bivalent promoters harbouring both marks are mainly poised and were observed in ES cells being associated to the pluripotent potential of these cells ^41^ and in differentiated cells where their roles remain to be addressed ^45,48^. We used H3K4me3 ChIP-seq and H3K27me3 Cut&Run to map these histone marks and again observed no differences between cells grown on stiff or soft hydrogels (Fig. 5a). However, we made two significant observations. First, the most differently expressed genes either on soft or on stiff condition display less H3K4me3 signals (Hi-soft and Hi-stiff found at top and bottom parts of the heatmaps, respectively, Extended Data Table 3), suggesting that responsiveness to mechano-transduction is compatible with lower levels of H3K4me3. This is consistent with the associated role of this mark to transcriptional consistency in particular at broad domains ^40^. Second, we found that genes that are more differentially expressed in soft (Hi-soft, top of the heatmaps) are enriched for both H3K4me3 and H3K27me3 histone marks, suggesting bivalent marking of the chromatin of these promoters. These genes are repressed under stiff ECM as expected but show significant expression levels when cells are grown under soft gels (Fig. 5a-b). We confirmed this tendency at the genome-wide level, where genes harbouring apparent chromatin bivalency are more expressed in soft as compared to stiff (Extended Figure 3a, Extended Data Table 4). While H3K27me3 marking deposited by PRC2 is generally associated with transcription inhibition, cells subjected to soft ECM seem to escape such repression in the context of bivalent chromatin. This could figure a novel mechanism for gene expression regulation through PRC2-dependent repression. Furthermore, genes from this group are enriched in biological processes linked to cellular adaptation to ECM modifications such as signalling proteins and components or regulators of extracellular matrix organisation and focal adhesion (Extended Data Fig. 3b).

**Figure 5:**
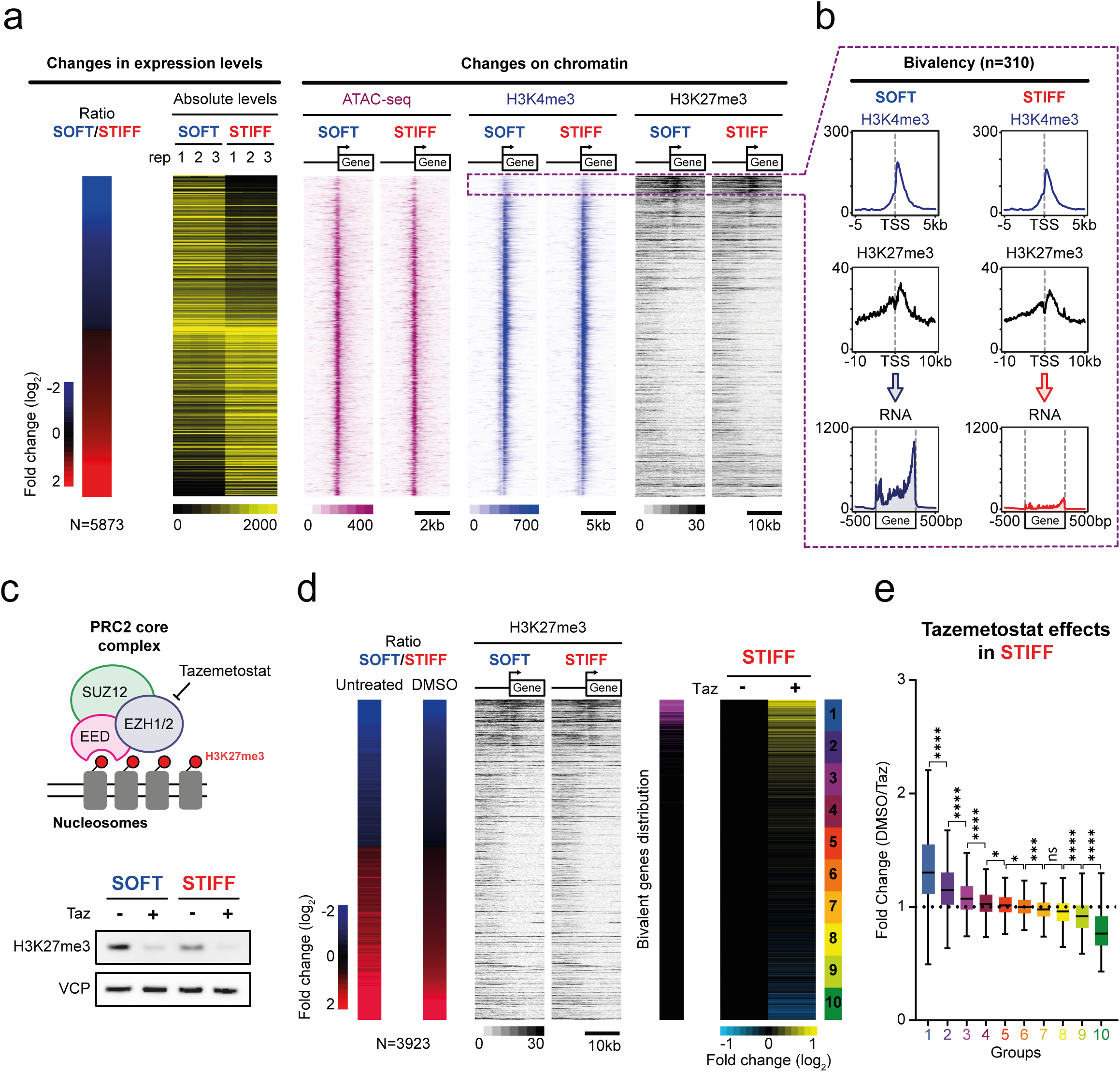
The SOFT gene signature involves H3K27me3-dependent repression of apparent bivalent genes. a. Heatmaps showing changes in gene expression of differentially expressed genes and their associated chromatin landscape (ATAC-seq, H3K4me3 ChIP-seq and H3K27me3 CUT&RUN). Genes are ranked based on the total RNAfold change between SOFT and STIFF conditions. Absolute levels: absolute number of reads from DESeq for each gene. A dashed rectangle highlights a group of genes harbouring both H3K4me3 and H3K27me3. b. Metaprofiles confirming the apparent chromatin bivalency and the changes in RNA levels of genes identified in **a**. c. Top: schematic representation of the core PRC2 complex and mode of action of the Tazemetostat drug. Bottom: western-blot comparing H3K27me3 signals without (-) and with (+) Tazemetostat treatment in each stiffness condition. VCP is used as loading control. d. Effects of Tazemetostat treatments on gene expression of mechano-responsive genes. Genes are ranked based on the fold change in expression between SOFT and STIFF hydrogels in DMSO condition. Fold changes in total RNA and H3K27me3 signals in untreated condition are also displayed. Pink: distribution of apparent bivalent genes identified in **a** and **b**. Right: Tazemetostat effect on gene expression in STIFF condition after Tazemetostat treatment (+) compared to DMSO (-). e. Tukey plots comparing fold changes of each of the 10 groups of genes determined in **d**. Statistical tests are Mann-Whitney.

To test the role of PRC2 in the inhibition of these set of genes, we used Tazemetostat, an FDA-approved EZH2 inhibitor used in the clinic for cancer treatment ^65,66^. Primary fibroblasts were treated for 7 days with 5µM of Tazemetostat resulting in 80 to 90 % decrease in H3K27me3 signal (Fig. 5c, Extended Data Fig. 4a). We then used total RNA-seq and focused on genes behaving similarly in untreated and control condition since adding DMSO has only slight effects on gene expression (Extended Data 4b, Extended Data Table 5). We found that under stiff condition the genes that display both H3K4me3 and H3K27me3 are indeed derepressed by Tazemetostat treatment (Fig. 5d). We also note that Hi-stiff genes are also down regulated in soft condition suggesting an indirect or non-canonical role of EZH2, independent of H3K27me3 marking, in the regulation of stiff-induced genes (Extended Data Fig. 4c-d).These results are consistent with our signature analysis that highlights the overlap of mechano-responsive genes with genes that were de-repressed in our soft signature or down-regulated in our stiff signature in fibroblast with EZH2 knock-down ^60^. Our genomic and functional analyses show that a set of Hi-soft genes with higher expression under soft ECM are marked in a bulk population by both H3K4me3 and H3K27me3 and require functional PRC2 for an efficient repression under stiff ECM.

### The nuclear lamina integrity is required for proper PRC2-dependent repression of genes associated to bivalency

As cells adhere and pull on their surrounding environment, the nuclear lamina is changing ^67^. In our primary fibroblast model, ECM stiffness promotes lamina component rearrangements, impact chromatin polymer properties and 3D genome modifications. To further address the NL role in softness-dependent and PRC2-dependent transcription programs, we compared our Hi-soft signature to LADs mapped by LaminB1 Cut&Run. Lamin B1 is part of the lamina meshwork facing the inner part of the nucleus (Fig. 6a). We first confirmed that globally, genomic regions covered by broad H3K27me3 domains are more associated to Lamin B1 than inter-LAD regions (Extended Data Fig. 5a). As exampled by the *BACH*2 gene, embedded in a H3K27me3 domain and associated to a large Lamin B1 domain (Fig. 6b), genes of the Hi-soft signature tend to be more associated to the NL than the rest of the genome or other soft-responding genes (Fig. 6c). This is observed in cells plated on both soft and stiff hydrogels since LAD domains remain largely unaffected by matrix stiffness. To further investigate if NL and the LINC complex are required for mechano-dependent transcription programs, we used siRNA targeting main NL components (LBR (LBR), Lamin A (LMNA), Lamin B1 (LMNB1) and Emerin (EMD)) and SUN proteins (SUN1, SUN2 and SUN1/2 in combination). We set scrambled siRNA as control (siNeg) in these series of experiments since transfection had some effects on the cellular response to ECM. We then performed total RNA-seq and only considered genes responding similarly to ECM stiffness with and without transfection (Extended Data Fig. 5b, Extended Data Table 6). We first confirmed the reduced RNA levels of our targeted genes (Extended Data Fig. 5c) and measure the effects of these treatments on the expression of the mechano-sensitive genes. Strikingly, we found that our Hi-soft signature is escaping repression on stiff condition when the NL is impaired, in particular when using siRNA targeting the LBR, LMNA or LMNB1 (Fig. 6d-e). We also noticed that siRNA treatments only have slight effects in soft condition suggesting that NL regulates gene expression mainly in response to stiff condition (Extended Data Fig. 5d-e). Hence, the NL integrity is required for proper PRC2 inhibition of the Hi-soft gene signature in stiff condition. To conclude, we have defined a Hi-soft gene signature enriched for mechanical-adaptive genes in our gene ontology analysis, controlling processes of ECM regulation, cell adhesion and signalling pathways. This set of genes is found associated to Lamin B1 in both condition but only expressed in cells platted on soft hydrogels. Under this condition, the NL is weakened, the chromatin is highly mobile and long-range 3D interactions are favoured. Under stiff condition, repression of these genes is altered when the NL is targeted by siRNAs or when EZH2 is inhibited by Tazemetostat. This supports the idea that thick and unaltered NL is required for PRC2-dependent Hi-soft gene repression. On soft hydrogels, wrinkled nuclei with loose lamina and mobile chromatin permit expression of this specific set of genes. We thus propose a novel regulatory mechanism where modifications in nuclear mechanics modulate chromatin mobility and 3D genome organisation. Under stiff ECM, nuclear flattening and stiffening associated with NL thickening create a constrain and rigid environment where chromatin motion and long-range 3D genome interactions are limited. This represents conditions for efficient PRC2 repression of genes with bivalently marked chromatin with both H3K4me3 and H3K27me3 histone modifications. Repression that could be tempered using the EZH2 inhibitor Tazemetostat or by targeting the NL components (Fig. 7).

**Figure 6:**
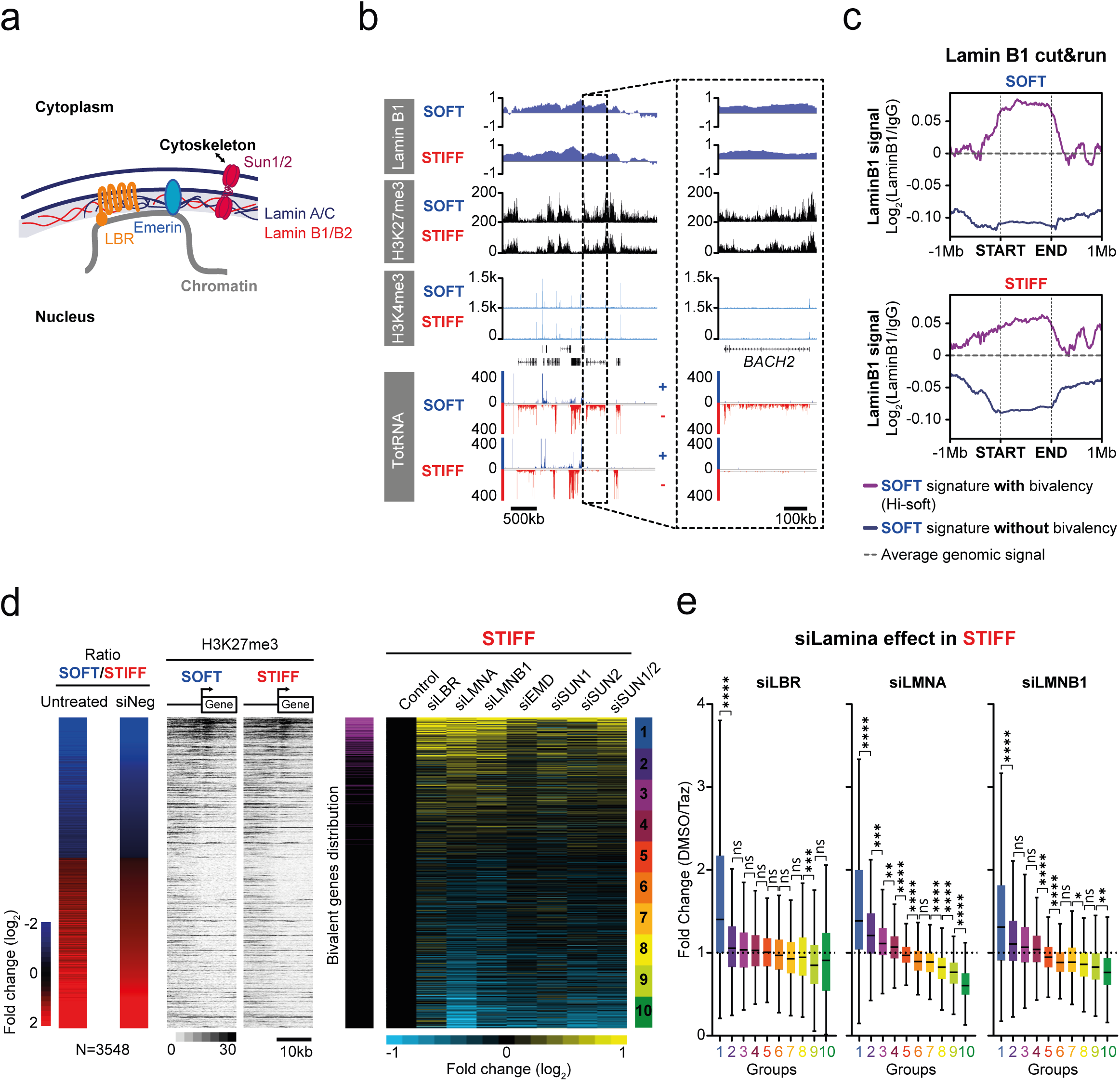
The nuclear lamina integrity is required for a proper repression of apparent bivalent genes in STIFF condition. a. NL scheme illustrating NL (Lamin-A/C and Lamin B1) and NL associated proteins (LBR and Emerin) as well as SUN proteins. b. IGB screenshots of Lamin B1 (light blue), H3K27me3 (black), H3K4me3 (cyan) and total RNA (blue for sense and red for anti-sense) signals on a genomic region containing the *BACH2* gene. c. Metaprofiles of Lamin B1 signals on apparent bivalent genes (Hi-soft, purple) and the other genes more expressed on SOFT hydrogels (blue). The dashed black line representes the average LaminB1 signal on the genome. d. Effects of siRNA treatments on gene expression of mechano-responsive genes. Genes are ranked based on the total-RNA fold change between SOFT and STIFF hydrogels in siNeg condition. Fold changes in total-RNA and H3K27me3 signals in untreated condition are also displayed. Pink: distribution of apparent bivalent genes. Right: siRNA targeting NL or NL-associated proteins effects on gene expression in STIFF condition (+) compared to siNeg (-). e. Tukey plots comparing fold changes of each of the 10 groups of genes determined in d for siLBR, siLMNA and siLMNB1. Statistical tests are Mann Whitney.

**Figure 7:**
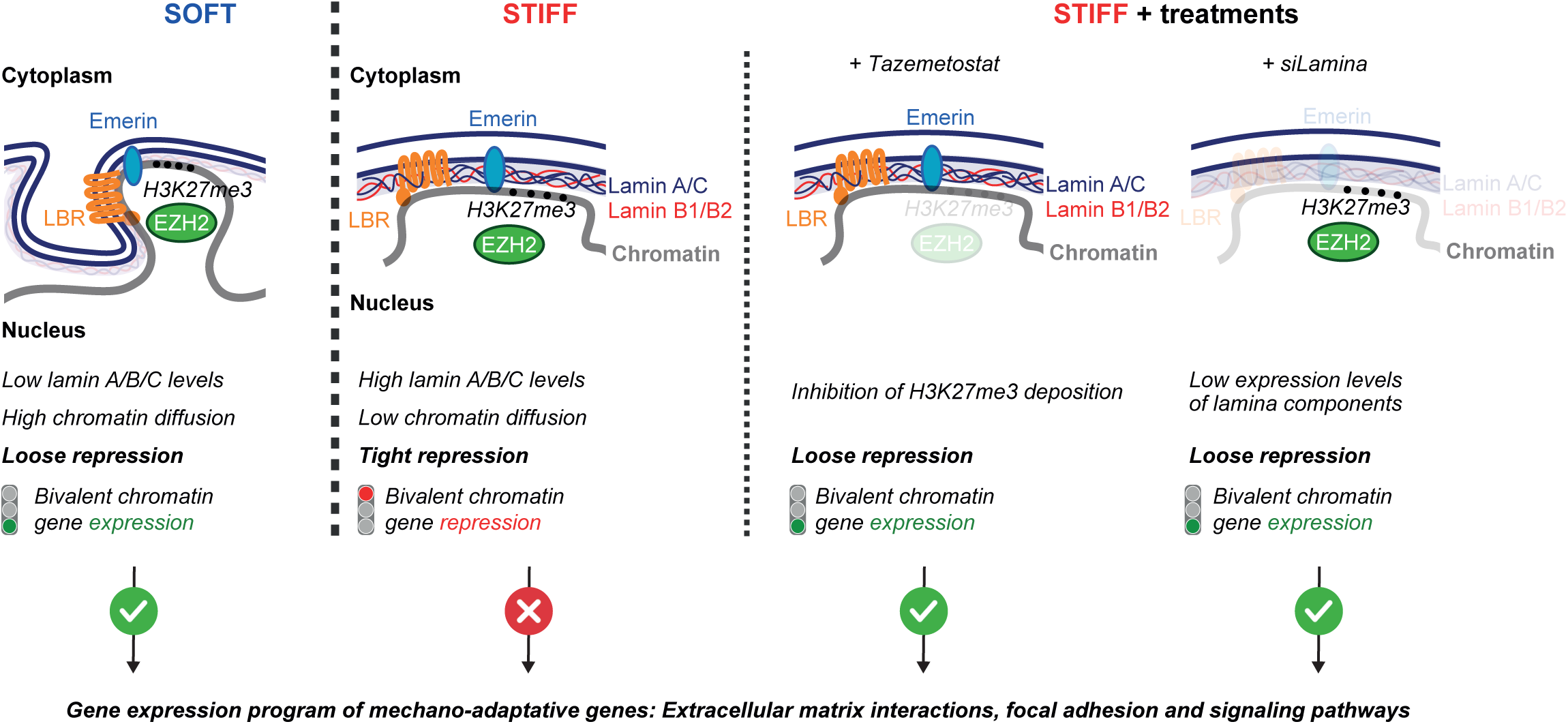
Model of bivalently marked genes regulation by mechanical cues. In both stiffness conditions, we identified genes marked by both H3K4me3 and H3K27me3 and associated to NL. In SOFT condition, Lamin A/C and Lamin B1 are lowly expressed, leading to a loose and wrinkled NL. This low physically constrained context allows chromatin motion, favour 3D genomic interactions and bivalent genes expression. In STIFF condition, NL is reinforced with high expression of Lamin A/C and Lamin B1 proteins. The nuclear flattening and NL stiffening creates a highly rigid and restrictive environment inducing the reduction of chromatin diffusion and the repression of bivalent genes. Tazemetostat targeting EZH2 or siRNA targeting NL main components lead to the re-expression of bivalent genes in STIFF condition, confirming that both PRC2 and the NL are required for the proper repression of these bivalently marked genes in this condition. Finally, this set of gene code for mechano-adaptive genes regulating extracellular matrix interactions, focal adhesion and signalling pathways.

## Discussion

Our data show that the NL and effective gene repression by PRC2 are mechano-sensitive and regulate the expression of specific sets of genes harbouring both H3K4me3 and H3K27me3 histone modifications. These genes are enriched in effectors of ECM adhesion and signalling. Antagonistic histone modification signatures that combine the activating histone H3K4me3 and the repressive H3K27me3 marks have been defined as bivalent. Bivalent marking was first identified in mES cells at genes characteristic of differentiation and developmental processes ^43–47^. While bivalency is proposed to contribute to the flexibility of the activation and repression of differentiation genes, its occurrence is not restricted to pluripotent cells and has also been observed in non-pluripotent cells ^45^. Furthermore, bivalent domains of terminal stages of differentiation can be distinct to those of ES cells ^48^ and their function remains quite unclear. We propose that they are at play in an adaptative mechanism where the NL acts as a sensor of extracellular matrix stiffness. As the environmental elasticity conditions are changing, the focal adhesion foci and the cytoskeleton adapt, resulting in profound cellular modifications. The NL organisation is in turn changing and regulates nuclear physical properties. This finally tunes the repression of genes harbouring bivalent marking of H3K4me3 and H3K27me3. In this model, bivalency would allow plasticity in the repression of mechano-adaptative genes which would thus respond to the modifications of the physical properties of the nucleus. Due to the limitation in primary fibroblast cell culturing, re-ChIP or similar approaches could not be performed and thus an alternative model would be that at the bulk population level, this specific set of gene would be differently marked at the subpopulation scale with a group of cells harbouring the H3K4me3 marking and another H3K27me3. This model remains however unlikely since only gene expression is changing without affecting the balance of H3K4me3 and H3K27me3.

This mechanism may also act during differentiation since the NL evolves during this process. The balance between Lamins A/C and the LBR changes with high levels of LBR and low levels of Lamins A/C in ES cells and with the opposite at the latest stages ^68^. This evolution is very similar to the differences we observed in response to ECM stiffness and may thus contribute to changes in expression and further modifications in the epigenetic marking of the bivalent genes. Similarly, our model could also contribute to aging since recent works have identified alterations in extracellular matrix composition and mechanics during aging of murine hair follicle stem cells ^9^. In this model, ECM is stiffening with aging concomitantly with the loss of bivalency with decreasing H3K4me3 signal and chromatin opening at bivalent genes.

Associations of H3K27me3 domains and LADs are still in debates. LADs are enriched in repressive marks including H3K27me3 ^27,30,32^ which may contribute to the attachment of the chromatin to the lamina through its interplay with YY1 ^31^. Moreover, Lamin-A/C was shown to interact with PRC2 proteins regulating their foci architecture and PRC2-dependent repression of myogenic genes ^69^. In contrast, recent works propose that H3K27me3 and PRC2 could antagonise LADs ^70,71^. However, one of these studies was conducted in the non-adherent cancer cell line K562 ^71^, while the other focused on early mouse embryonic development, a context in which non-canonical H3K27me3 domains have been described ^70,72^. These differences in cellular context, developmental stage, mechanical states and probably in NL composition may underlie the discrepancies between our findings and those previously reported.

Our gene expression assays show that a very large set of genes is mechano-responsive. Our ontology analysis indicates more house-keeping gene functions being enhanced under stiff condition. Due to the limited duration of our experimental setup, BJ primary fibroblasts which have a doubling time of over a week under hydrogel were not afforded sufficient time to initiate or complete cell proliferation. However, our assays predict that long term exposure on differential hydrogel stiffness would impact cell fitness and proliferation. In addition, our assays also highlight the transcriptional regulation of TNF, AKT or MAPK signalling components in the soft responding gene signature. These signalling pathways are proposed to be at play in response to mechanical inputs or focal adhesion dependent signalling ^73–75^. Hence, further investigation of these signalling cascade dependencies and activations in response to ECM stiffness and to its gene expression program regulation would be of great interest. In particular for AKT dependent signalling since it controls both NFKB and Foxo transcription programs ^76,77^ for which binding motif enrichments were also identified in our study.

3D genome organisation and gene regulation are paired. Here, we found that long range interactions at the compartment scale are influenced by the ECM stiffness. Under soft growth condition, the frequency of interactions at this scale is increased. This also correlates with LADs and with changes in chromatin motion. Chromatin in the nucleus can be model and its viscoelastic displacement are consistent with Rouse polymer models ^78^. Our HiD measurement indicates directed motion in cells grown on soft hydrogels close to the nuclear periphery that would be consistent with broad movement of the NL and of its associated chromatin domains. These movements could also represent a novel layer of gene regulation by the NL and its flexibility. Increase in motion of large chromatin domains linked to the NL could thus enhance the likelihood of long-range interactions measured by HiC and loose repression that occurs at sub diffusing domains in stiff condition. We propose that flattening of the nucleus associated to NL thickening change nuclear mechanical properties, 3D genome organisation as well as chromatin motion and promote H3K27me3-dependent repression of genes with bivalent chromatin found close to NL by creating a more restrictive environment.

All in all, our data suggest that NL acts as a mechano-sensor and effector to control specific sets of adaptative genes that are bivalently marked. We propose a gene expression control mechanism where modifications in the NL organisation in response to extracellular matrix stiffness leads to nuclear and chromatin mechanics modifications. These changes would set conditions for efficient or loose PRC2-dependent repression of bivalently marked genes. Hence, sensed mechanical inputs through the NL would control a specific set of genes regulating biological pathways involved in the cellular adaptation to mechanical cues. These findings could be further used to understand how 3D genome rewiring could promote mechano-dependent physiological effects such as tumour progression and stem cell differentiation and to define PRC2 as a potential target for treatments..

## Experimental procedures

### Antibodies and applications

All antibodies and their applications are listed in Extended Data Table 7.

### Cell amplification

Primary BJ fibroblasts were expanded on plastic plates in DMEM High Glucose (ThermoFisher, 31966021) supplemented with 10% foetal bovine serum (FBS, Gibco) at 37°C and 5% CO2.

### Hydrogel-based cell culture

Polyacrylamide hydrogels (Matrigen, Cell Guidance System) with a stiffness of 0.5kPa (SOFT) and 50kPa (STIFF) were coated with fibronectin from human plasma (Sigma, F0895) at a final concentration of 10µg/mL for 2h at room temperature (RT). Then, cells were trypsinised and plated on fibronectin-coated hydrogels in DMEM High Glucose (ThermoFisher, 31966021) supplemented with 10% foetal bovine serum (FBS, Gibco) at 37°C and 5% CO2 for 48h before performing the subsequent experiments. Different types of hydrogels were used depending of the experiment. For RNA and proteins isolations: Softwell 6-well plate Easy Coat (SW6-EC-0.5 and SW6-EC-50), for microscopy experiments: SoftSlip 12-well plate Easy Coat (SS12-EC-0.5 and SS12-EC-50), for genomic analysis requiring large amount of cells: PetriSoft 150mm Dish Easy Coat (PS150-EC-0.5 and PS150-EC-50) and for live microscopy HiD experiments: Softview 35mm dishes (SV3520-EC-0.5 and SV3520-EC-50).

### siRNA and Tazemetostat treatments

#### siRNA

Cells were transfected using Lipofectamin RNAiMAX (Invitrogen, 13778150) and 20nM of siRNA (Dharmacon, Horizon Discovery) according to the protocol of reverse transfection to maximise the efficiency. All siRNA references are listed in Extended Data Table 8. Briefly, for one 10cm dish, 15µL of Lipofectamin RNAiMax was diluted in 1mL of Opti-MEM medium (Gibco, 31985062) and incubated for 5min at RT. Simultaneously, siRNA were diluted in 1mL of Opti-MEM medium and incubated 5min at RT. Both solutions were then mixed at a 1:1 ratio into a 10cm dish and incubated 20min at RT. Resuspended trypsinized cells were added for a final concentration of siRNA at 20nM and incubated 3 days in DMEM High Glucose 10% FBS at 37°C and 5% CO2. Afterwards, cells were trypsinised and plated on fibronectin-coated hydrogels for 48h with 20nM siRNA.

#### Tazemetostat

Cells were plated on plastic plate and treated for 5 days with 5µM of Tazemetostat (Selleckchem, EPZ-6438) in DMEM High Glucose 10% FBS at 37°C and 5% CO2. Media was changed each 2days. Afterwards, cells were trypsinized and plated on fibronectin-coated Softwell 6-well plate Easy Coat hydrogels (SW6-EC-0.5 and SW6-EC-50) for 48h with 5µM Tazemetostat.

### Confocal and super-resolution microscopy

Cells were fixed directly in the wells with freshly prepared 4% paraformaldehyde in PBS 1X for 10min at RT, permeabilised with 0.2% Triton X-100 for 10min at RT and incubated in blocking solution (10% FBS, 1% Cold Fish Skin Gelatin in PBS 1X) for 10min at RT. Primary antibodies were diluted in blocking solution and incubated for 2h at RT. Secondary antibodies as well as phalloidin labelling actin filaments were diluted in blocking solution and incubated for 1h at RT. Nuclei were stained with DAPI for 10min at RT. After extensive washes with PBS-Tween 0.5%, a final wash with H2O was performed to remove salts. A drop of Prolong Gold (Invitrogen, P36930) was deposited on the microscopy slide to mount the coverslip containing-hydrogel, 10µL of Vectashield was used to mount the coverslip on the hydrogel and transparent nail polish was immediately used to seal the mounting. Confocal images were acquired with a Zeiss LSM780 and super-resolution images with a DeltaVision OMX 3D-SIM.

### Live microscopy for HiD

Three days prior imaging, cells were transfected with H2B-GFP with FuGene HD transfection reagent (promega, E2311) at a ratio of 4:1. Two days before, cells were transferred in Softview dishes of 0.5 and 50kPa respectively. On the day of imaging, culture medium was changed to L-15 medium (gibco, 21083-027) supplemented with 10% FBS. Cells were placed in a 37°C, 5% CO2 humid chamber (OkoLab system). Imaging was performed using a DMi8 inverted automated microscope (Leica Microsystems) featuring a confocal spinning disk unit (CSU-X1-M1N, Yokogawa). A 63x oil immersion objective (Leica HC-PL APO, 11506349) with a 1.4 NA was used for high resolution imaging. Image (16bit) was acquired using Metamorph 7.10.5 software (Molecular Devices) and detected using a CMOS Hamamatsu Flash4 V2+ camera (1003x1433) and 1 x 1 binning, with sample pixel size of 120 nm. Image series of H2B-GFP (488 nm laser with a band pass emission filter of 510-540) of 500 frames, with exposure time of 200 ms per frame (5fps) were acquired.

### Western-blot

Cells were lysed directly in the well with RIPA buffer (10mM Tris-HCl pH 8.0, 1mM EDTA, 0.5mM EGTA, 1% Triton X-100, 0.1% Sodium deoxycholate, 0.1% SDS, 140mM NaCl). Protein extracts were then complemented with Laemmli 4X (250mM Tris ph6.8, 8% SDS, 40% glycero, 5% β-mercaptoethanol, bromophenol blue) to a final concentration of 2X, incubated 5min at 95°C on a thermomixer before loading on a 4-12% Bis-Tris gel. After the transfer on a nitrocellulose membrane, blocking solution (5% milk in PBS-Tween 0.2%) was added for 1h at RT. Primary antibodies were diluted in 2% milk PBS-Tween 0.2% and incubated O/N at 4°C. Secondary antibodies were diluted in 2% milk PBS-Tween 0.2%, incubated 1h at RT and revelation was performed with ECL (BioRad) or SuperSignal West Femto (Thermofisher) on an Amersham Imager 680 (GE LifeSciences). Bands quantifications were performed using the ImageJ software.

### Total RNA-seq

Cells were plated on Softwell 6-well plate Easy Coat hydrogels with each well representing one biological replicate. Total RNAs were purified using RNeasy Plus Mini Kit (Qiagen) and eluted in 30µL of nuclease-free water. Quantifications were performed using Nanodrop and qualities were assessed with a BioAnalyzer Pico RNA chip (Agilent). A fraction of purified RNA was used for subsequent reverse-transcription with the Superscript Vilo kit (Thermofisher, 11754250) and expression of known mechano-responsive genes such as *ANKRD1* and *CTGF* was assessed by qPCR. For RNA isolated from cells treated with siRNA, qPCR were performed on the targeted associated gene (all primers listed in Extended Data Table 9). Then, 10µL of total RNA was mixed with ERCC92 spike-in (Invitrogen, 4456740) depending on RNA quantity according to manufacturer instructions and treated with DNase before proceeding to libraries preparation using the TruSeq Stranded Total RNA Ribo-Zero H/M/R Gold (Illumina). Libraries were quantified using the Qubit dsDNA kit (Thermofisher) and size distribution was assessed with a BioAnalyzer High Sensitivity DNA chip (Agilent). All the fragments with a size superior to 700bp were cleaned with CleanNGS beads (CleanNA) and libraries were sequenced in 2x50bp paired-end on an HiSeq4000 or a NovaSeq.

### ATAC-seq

ATAC-seq experiments were performed as described previously in ^79^ in triplicates. Briefly, cells were trypsinised, washed in PBS 1X and counted in order to have ∼65000 cells per replicate. Cells were centrifugated 10min at 500g at 4°C and supernatant was discarded. Cell pellets were lysed with 50µL of cold lysis buffer (10mM Tris-HCl pH 7.5, NaCl 10mM, MgCl2 3mM, 0.1% Igepal CA-630). After centrifugation 15min at 500g 4°C, nuclei were resuspended into 50µL of tagmentation buffer (25µL of TD buffer, 2.5µL of Tn5 transposase, qsp 50µL with nuclease-free water) and incubated 30min at 37°C. DNA was then purified with MinElute columns (Qiagen) according to the protocol. Samples were indexed and amplified by PCR for 10 cycles to generate sequencing libraries. Libraries were purified using the PCR purification kit (Qiagen), quantified with Qubit dsDNA kit (Thermofisher) and deposited on a BioAnalyzer High Sensitivity DNA chip (Agilent). All fragments with a size superior to 300bp were eliminated with CleanNGS beads (CleanNA) to keep only the subnucleosomal fraction. Libraries were then sequenced on an HiSeq4000 in 2x75bp paired-end.

### ChIP-seq

ChIP-seq experiments were performed in triplicates. Approximately 8x10^5^ cells per replicate were collected by trypsinisation and washed with PBS 1X before fixation with 1% formaldehyde for 10min at 37°C. Reaction was stopped by the addition of glycine to reach a final concentration of 125mM and incubated 5min at RT. All centrifugation steps were done at 4°C 1200g. Cells were abundantly washed with cold PBS 1X, resuspended in 10mL of buffer A (5mM Hepes pH 7.5, 85mM KCl, 0.5% Triton X-100), incubated for 10min on ice. After centrifugation, 5mL of buffer A’ (5mM Hepes pH 7.5, 85mM KCl) were added. Cellular pellet was resuspended in 500µL of lysis buffer (50mM Tris-HCl pH 8.0, 1% SDS, 10mM EDTA, 1mM PMSF, 1X cOmplete Protease Inhibitor), incubated on ice for 10min before snap-freezing in liquid nitrogen. After defrosting, chromatin was sonicated with a Bioruptor Pico (Diagenode) for 4 cycles 30sec ON/30sec OFF. A 10µL aliquot was added to 90µL of FA/SDS/like buffer, 25µL of elution buffer 5X (125mM Tris-HCi pH 7.5, 25mM EDTA, 2.5% SDS) and 5µL of proteinase K before incubation O/N at 65°C. Next day, DNA was cleaned using PCR Purification kit (Qiagen) and eluted in 30µL of nuclease-free water. Then, DNA was quantified using Qubit dsDNA kit and ran on a 1.5% agarose gel to verify that mean DNA shearing was below 500bp. After validation of sonication profiles, samples were centrifugated to pellet cell debris, supernatant was recovered and diluted 10 times in FA/SDS/Like buffer (50mM Hepes pH 7.5, 150mM NaCl, 1% Triton X-100, 0.1% Na-Deoxycholate). Diluted chromatin was aliquoted in 1mL, snapped-freeze in liquid nitrogen and conserved at -80°C until use. Protein-G coated dynabeads (Thermofisher) were blocked with PBS-BSA 0.1% and 25µL of beads per immunoprecipitation (IP) was coupled with the antibodies of interest on a wheel for 2h at 4°C. An aliquot of 1mL of sonicated chromatin was precleared by addition of 25µL of dynabeads resuspended in PBS-BAS 1% per sample to reach a final concentration of 01% BSA and incubated on a wheel for 2h at 4°C. An aliquot of 20µL was taken as input and treated as described above for sonication check. Beads incubated with the antibodies were collected and washed 4 times with PBS-BSA 0.1% before addition of pre-cleared chromatin and incubation O/N at 4°C on a rotating wheel. The following day, beads were washed one time with 500µL of FA/SDS/PMSF buffer (50mM Hepes pH 7.5, 150mM NaCl, 1mM EDTA, 1% Triton X-100, 0.1% Na-Deoxycholate, 0.1% SDS) and 3 times with 500µL of FA/SDS/PMSF buffer 300mM NaCl. During the last wash, samples were placed on a rotating wheel 10min at 4°C. Then, beads were washed with 500µL of washing buffer (10mM Tris-HCl pH 8.0, 250mM LiCl, 1mM EDTA, 0.5% NP-40, 0.5% Na-Deoxycholate) and 500µL of TE buffer (10mM Tris-HCl pH 8.0, 1mM EDTA) before incubation 25min at 65°C in a thermomixer with 125µL of pronase buffer 1X (25mM Tris pH 7.5, 5mM EDTA, 0.5% SDS). Supernatant containing immunoprecipitated fragments was recovered, purified with PCR purification kit (Qiagen), eluted in 30µL of nuclease-free water and quantified using Qubit dsDNA kit (Thermofisher). Enrichment on target control regions was assessed by qPCR (primers listed in Extended Data Table 9) and sequencing libraries were prepared with MicroPlex Library Preparation V2 or V3 kits (Diagenode). Libraries were quantified using the Qubit dsDNA kit (Thermofisher) and size distribution was assessed with a BioAnalyzer High Sensitivity DNA chip (Agilent). All the fragments with a size superior to 700bp were cleaned with CleanNGS beads (CleanNA) and libraries were sequenced in 2x50bp paired-end on a NovaSeq.

### Cut&Run

Cut&Run experiments were performed in triplicates as previously described in ^80^ with minor modifications. Each Cut&Run was performed with 8x10^5^ cells. After cells permeabilisation with 0.05% digitonin and incubation with washed and activated beads, the mix was resuspended in 300µL of antibody buffer containing antibodies targeting either LaminB1, H3K27me3 or IgG (negative control) at a 1/100 dilution and incubated 2h at 4°C under agitation. After washing, samples were resuspended in 100µL of cold digitonin buffer and 5µL of diluted 1/10 pAG-MNase was added to reach a final concentration of 700ng/mL and incubated 10min at RT. After washing to remove unbound pAG-MNase, 100µL of cold digitonin buffer was added as well as 2µL of CaCl_2_ 100mM. Samples were incubated 2h at 4°C under agitation to proceed to the digestion process. Reaction was stopped by the addition of 66µL of Stop Buffer 2X and digested chromatin was released in the supernatant by incubation 10min at 37°C. Supernatant was recovered and purified with Monarch kit (NEB). Sequencing libraries were prepared using the MicroPlex Library Preparation V3 kit (Diagenode), quantified with Qubit dsDNA, and size distribution was assessed with a BioAnalyzer High Sensitivity DNA chip (Agilent). All the fragments with a size superior to 700bp were cleaned with CleanNGS beads (CleanNA) and libraries were sequenced in 2x50bp paired-end on an HiSeq 4000 or a NovaSeq.

### Hi-C experiments

Hi-C experiments were performed in duplicates with the Arima Hi-C kit (Arima Genomics, Inc.; Cat # A510008) according to the manufacturer instructions. One million cells were crosslinked with formaldehyde and subjected to Arima HiC Standard Input protocol. Briefly, cells were lysed and chromatin was digested using a combination of restriction enzymes. The 5’-digested ends were filled in with a biotinylated nucleotide and spatially proximal digested ends were ligated. Proximally ligated DNA was purified using CleanNGS beads (CleanNA) and fragmented using the Covaris sonicator to reach an average size of 400bp (temperature: 20°C ; peak incident power: 50W ; duty factor: 10% ; cycles per burst: 200cpb ; time: 70s). Fragments between 200bp and 600bp were size-selected and biotinylated ones were enriched using furnished streptavidin enrichment beads. On-beads sequencing libraries were prepared using the MicroPlex Library Preparation V2 kit (Diagenode), quantified with Qubit dsDNA and size distribution was assessed with a BioAnalyzer High Sensitivity DNA chip (Agilent). All the fragments with a size superior to 700bp were cleaned with CleanNGS beads (CleanNA) and libraries were sequenced in 2x50bp paired-end on an HiSeq 4000.

### Bioinformatic procedures

#### Quantifications of nuclear volume and nuclear lamina thickness

Nuclear volumes and NL thicknesses were quantified using ThicknessGui (https://github.com/ant-trullo/ThicknessGui). Briefly, after opening the Z-stack files obtained from super-resolution acquisitions, we manually selected the frames of interest and divided our nuclei in 180 slices. After calculation, we exported the .csv file and the visual representation of nuclei segmentation. We then selected the slices that fit the best with our Lamin A/C signal. Since the values in the .csv files are pixels, we multiplied by pixel size (pixel width: 0.4µm, height: 0.4µm, depth: 0.125µm) to obtain NL thickness and nuclear volume values in µm and µm3, respectively.

#### HiD data processing

Before HiD analysis, nuclei with a low GFP intensity (less than two time the intensity of the background) and which undergo Z-plan drift were eliminated. Bleaching and XY-plan drift were corrected using Bleach Correction (<u>10.12688/f1000research.27171.1</u>) and StackReg translation correction (<u>10.1109/83.650848</u>) imageJ plugins respectively. Finally, to remove high gain noise from time laps image streams, the Kalman Stack Filter plugin from image J is applied.

HiD analysis is based on the method of (10.1186/s13059-020-02002-6, 10.1038/s41596-024-01038-3). Briefly, detection of movement between two consecutive frames were computed by the Farneback Optical Flow based on polynomial expansion (10.1007/3-540-45103-X_50). For a sequence with N frames, N-1 flow fields were produced and linked to compute a trajectory per pixel. The MSD curves are computed for every trajectory as function of the time interval (τ)and the first 20% (100 frames = 20s) of the experimental MSDs were fitted with the following anomalous diffusion model:

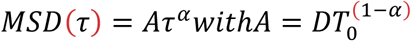

Where A (in µm^2^/s^α^) is the product of the diffusion constant D (in µm^2^/s) and T0 (in s), a characteristic time at which a diffusive particle sense its neighbours. α is its anomalous exponent allowing to characterise three model of diffusion, i. free diffusion, α=1, ii. subdiffusion or anomalous diffusion with 0<α<1, iii. superdiffusion or directed motion with 1<α.

#### RNA-seq data processing

Raw sequencing reads were aligned to the human genome (hg19) using TopHat2^81^. Sequences aligning on positive and negative strands were separated and processed separately using PASHA pipeline ^82^ to generate strand-specific WIG files representing the average enrichment score every 50bp. All WIG files were then rescaled to normalize the enrichment scores to the depth of sequencing. Rescaled WIG files of biological replicates from the same condition were then merged to produce merged WIG files. Reads Per Kilobase per Million (RPKM) were generated based on DESeq values normalised by annotation size.

#### ATAC-seq, ChIP-seq and Cut&Run data processing

Raw sequencing reads were aligned to the human genome (hg19) using Bowtie2^83^. WIG files were generated using the PASHA R pipeline ^82^. For ATAC-seq, WIG files represent average enrichment score over 10bp while for ChIP-seq data and Cut&Run targeting H3K27me3, those represent the average signal over 50bp and for Lamin B1 Cut&Run over 10kb. All WIG files were then rescaled to normalize the enrichment scores to the depth of sequencing. Rescaled WIG files of biological replicates from the same condition were then merged to produce merged WIG files. For Lamin B1 and negative control IgG files, signal over 10bins, representing signal on 100kb, was averaged to produce a rescaled and normalized WIG with a resolution of 100kb. Then, Lamin B1 signal over 100kb was normalized with negative IgG control and logged in log2 to highlight Lamin B1 associated domains.

#### H3K4me3 peak-calling

Peak-calling was performed using MACS2 ^84^. Only peaks found in at least 2 over 3 replicates are conserved. For ChIP-seq targeting H3K4me3, peak-calling was performed with the merged bam file of control IgG in 0.5kPa and 50kPa as background using the parameter -c. All peaks with a threshold of p-value < 10^-15^ were selected and conserved based on the “narrow peak” table provided by MACS2. DESeq package was used to quantify reads mapping on the called peaks and Reads Per Kilobase per Million (RPKM) were generated using DESeq output values normalised by domain size.

#### H3K27me3 domains calling

H3K27me3 bam files from the same stiffness condition were merged and used for peak-calling. Peak-calling as performed using MACS2 ^82^ with parameters --keep-dup all, -f BAMPE, -maxgap 3000. The merged bam file of control IgG in 0.5kPa and 50kPa was used as background with the parameter -c. All peaks with a threshold of p-value < 10^-3^ were selected and conserved based on the “narrow peak” table provided by MACS2. DESeq package was used to quantify reads mapping on the called domains and Reads Per Kilobase per Million (RPKM) were generated using DESeq output values normalised by domain size.

#### Lamin B1 domains calling

Lamin B1 WIG files normalized on the depth of sequencing were then normalized over IgG control signal and logged in log2 to allow domain visualization on the IGB tool. Then, wigs from the same stiffness condition were merged and peak-calling was performed using WigPeakCalling script ^63^ which extracts values from the thresholding function of IGB. The parameters used were: -- minRun 30000 and --maxGap 50000. DESeq package was used to quantify reads mapping on the called domains and Reads Per Kilobase per Million (RPKM) were generated using DESeq output values normalised by domain size.

#### Differential Expression Analysis

Differential Gene Expression (DGE) analysis was performed using the DESeq package ^85^. First, HTseq-count ^86^ was used to count the sequenced reads mapping to hg19 Refseq protein coding gene annotations or protein coding genes introns annotations. These counts were merged in order to have tables containing all biological replicates from different conditions. Then, these tables were processed using the DESeq package to identify genes that are significantly differentially expressed relative to the reference sample. A threshold of padj < 0.01 and Fold change < 1 (more expressed on SOFT) or > 1 (more expressed on STIFF) was used to identify differentially expressed genes in untreated samples and a pval < 0.05 was used for Tazemetostat and siRNA treated samples.

#### Coverage calculations

Coverage represents the number of base pairs covered by an epigenetic mark on an annotation of the genomic reference. H3K27me3 domains were characterised by peak calling and a bed file containing the corresponding coordinates was generated. This bed file was then intersected with interLAD or LAD annotations. All base pairs of an annotation covered by the epigenetic mark were accumulated and divided by the annotation size.

#### Gene ontology and gene signature enrichment

Gene ontology analysis was performed using DAVID software ^87,88^. Signature enrichment studies were completed using DESeq output data used as input in GSEA software ^89^.

#### Average binding profiles and heatmaps

To generate average binding profiles of Lamin B1, annotation files containing the 311 apparent bivalent genes in one hand and all genes more expressed on SOFT (identified with DESeq: padj < 0.01 and FC < 1) in another hand were generated. Log2-normalised Lamin B1 WIG files were converted into bigWIG files and used in the Deeptool software ^90^ to extract signal in a window of 1Mb around the selected annotations. Heatmaps were generated, viewed and colour-scaled using Java TreeView ^91^. Regions were centred on the annotated transcription start sites (TSS) of the differential protein coding genes identified and signal in defined windows was retrieved from WIG scaled files.

#### Identification of bivalent genes at genome-wide level

Promoter regions for all protein coding genes from RefSeq hg19 annotations were defined as a window of 300bp before and 100bp after the annotated transcription start site (TSS). Using the bedtools intersect tool, promoter regions overlapping in their whole with H3K27me3 (in H3K27me3 domains), H3K4me3 (Others) or both (Bivalency - In H3K27me3 domains) were identified.

#### Hi-C analysis

Sequences were analysed and aligned using the HiCUP/bowtie2 pipeline ^92^ with the by default parameters. Briefly, the human genome (hg19) was digested in silico with the restriction enzymes of the Arima option to create a reference genome. Sequences were then cut at the putative ligation site and all artefacts such as re-ligation of adjacent genomic fragments or PCR duplicates were removed. The remaining were aligned using bowtie2 on the digested reference genome. Homer was then used to generate the interaction matrix.

### 3D contacts modelling

3D genomic contacts were modelled with the ShRec3D algorithm using the default paramaters ^58,93^on chromosomes from 11 to 20. Each colour represents one chromosome.

## Acknowledgements

This work was supported by a grant from “Agence Nationale de la Recherche” (MecEpi, ANR-18-CE12-0019), la Fondation ARC (PJA-2018), La ligue contre le cancer, le cancéropôle GSO Emergence (N°2022-E07 / CANCÉROPÔLE GSO). We are grateful to Robert Feil for critical reading of the manuscript.

## Author contributions

AP and CE designed experiments; AP and MRM performed the experiments; CE, AP and AZEA analyzed genomic data. AT designed the software for lamina thickness determination from high resolution microcopy images. MRM, LC, MM and KB performed the HiD experiments and developed the associated analytical tools. CE conceived and supervised the project, suggested and interpreted experiment. JCA advised on the conception and the supervision of the project. AP and CE wrote the article. All authors reviewed the manuscript.

## Data availability

The GEO accessions for specific experiments related to this study are recorded under GSE303010, GSE303013, GSE303014, GSE303016 and GSE303035.

---

**Extended Data Figure 1:**
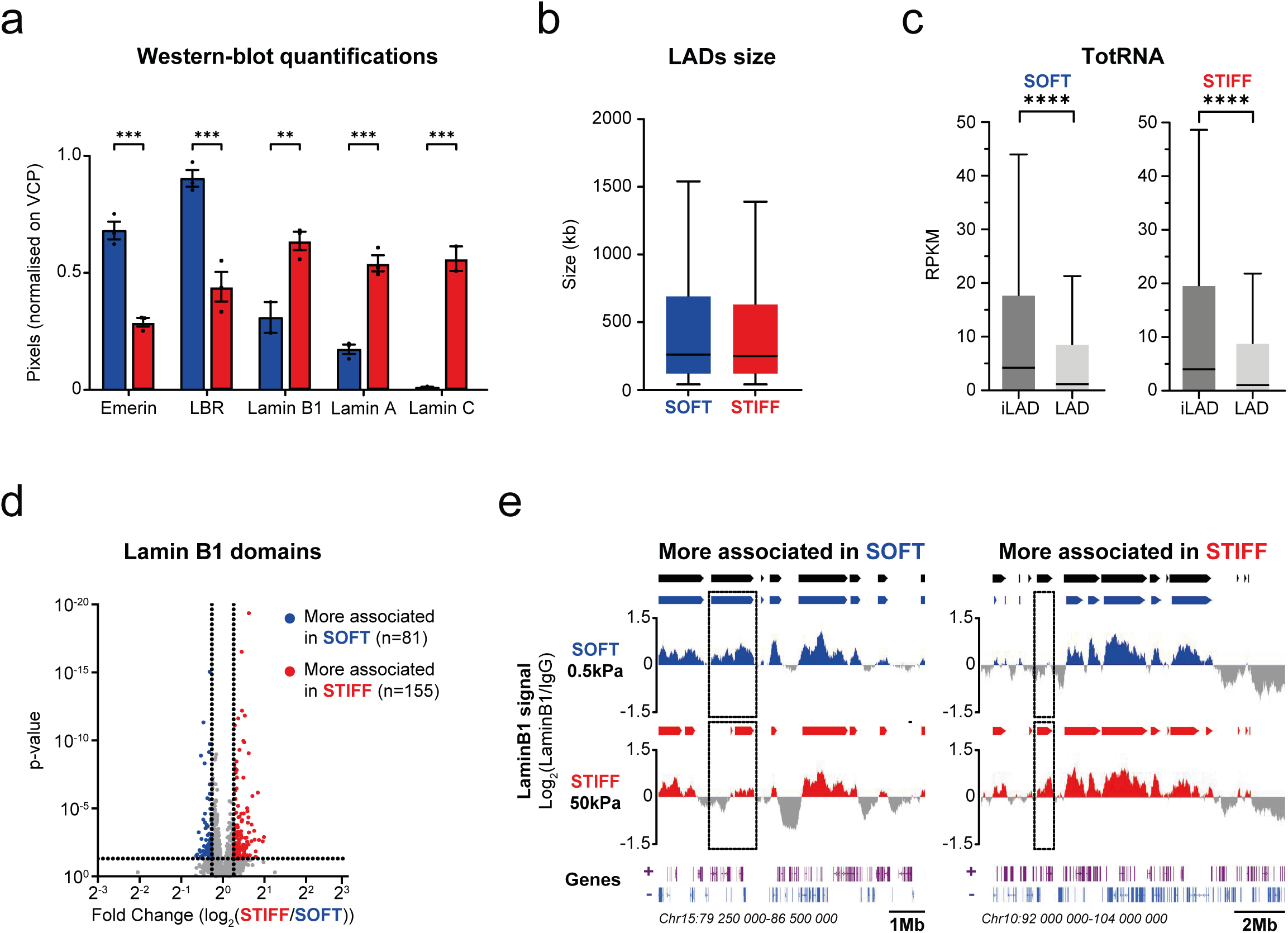
LADs organisation is largely unaffected by stiffness. a. Quantifications of NL and NL-associated proteins abundance by western-blot in Figure 1c. Mean with SEM of 3 independent replicates are represented. Statistical tests are unpaired two-tailed t-tests. b. Tukey plots analysing LADs size (in kb). Middle bar represents the median size. c. Tukey plots of Reads Per Kilobase per Million of reads (RPKM) of total RNAseq in interLAD (iLAD) and LAD regions for each stiffness condition. Middle bar represents the median. Statistical tests are Mann-Whitney. d. Volcano plot of genomic regions differentially associated to Lamin B1 between stiffness conditions (DESeq package; threshold: pval < 0.05, Fold-Change: 1.2). Genomic regions more associated on SOFT are in blue and those more associated on STIFF hydrogels are in red. e. IGB screenshots of genomic regions illustrating a genomic locus more associated to Lamin B1 in SOFT (left) or in STIFF (right) hydrogels. Arrows on top represent LADs called in SOFT (blue), in STIFF (red) or the merge of both conditions (black). Region more associated in SOFT: Fold-change = 0.69 and p-value = 1.3e^-9^. Region more associated in STIFF: Fold-change = 1.54 and p-value = 4.42e^-20^.

**Extended Data 2:**
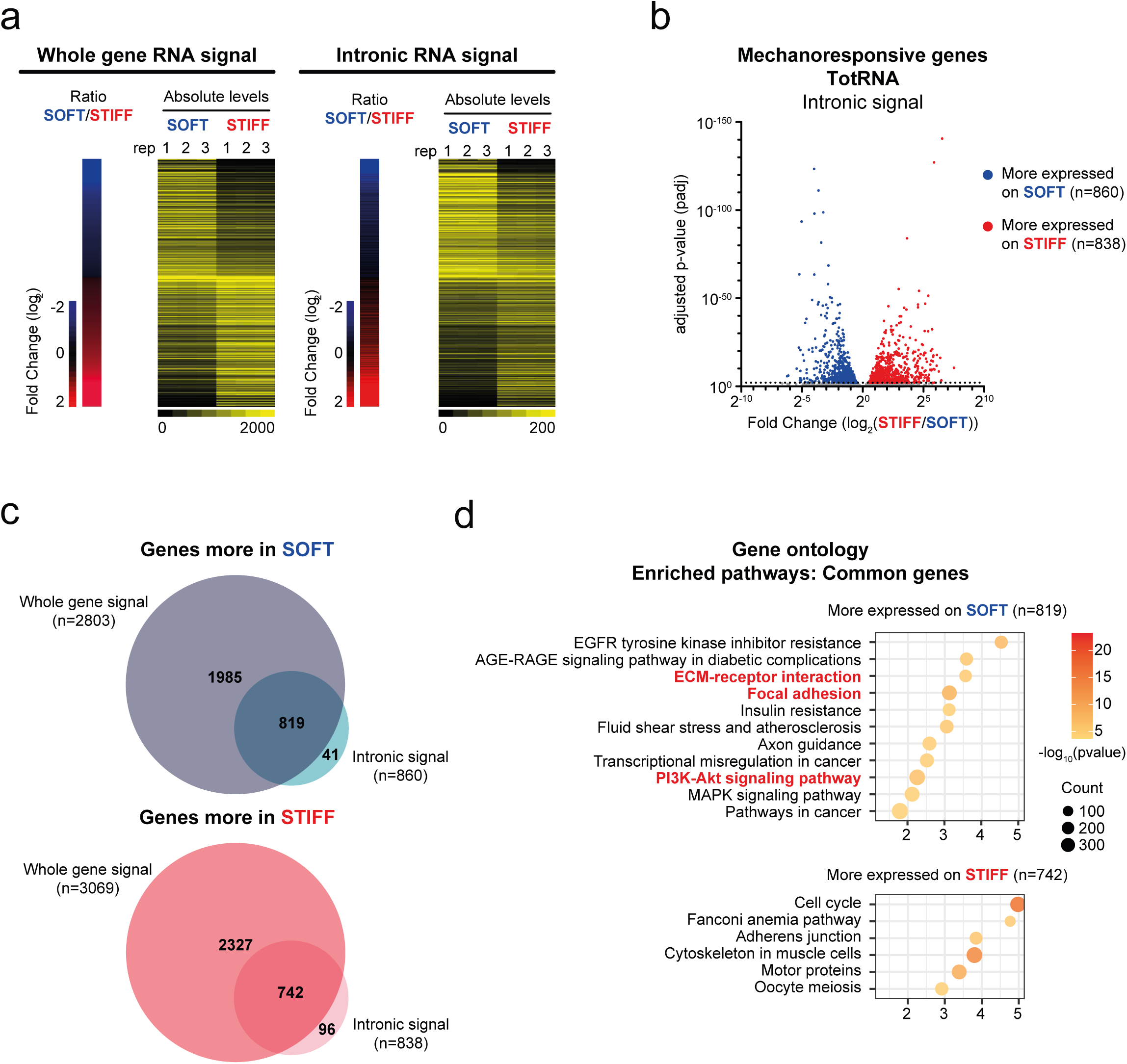
Matrix stiffness regulates gene expression programs at transcriptional levels. a. Comparison of gene expression changes driven by matrix stiffness at whole protein coding gene and intronic reads levels. Genes are ranked based on their RNA-seq fold change between SOFT vs STIFF hydrogels at the whole gene level. Associated ratio at intronic levels are shown on the right. Absolute levels of expression corresponding to the number of reads from DESeq are shown for both whole and intronic signals. b. Volcano plot of differentially expressed genes between soft and stiff hydrogels at intronic levels. ( DESeq package; threshold: padj < 0.01, Fold-Change = 1). Genes more expressed on SOFT are in blue and those more expressed on STIFF hydrogels are in red. c. Venn-diagrams illustrating the genes that are differential at whole, intronic or both levels depending on stiffness conditions. d. Gene ontology (GO) analysis of differentially expressed genes at intronic levels.

**Extended Data 3:**
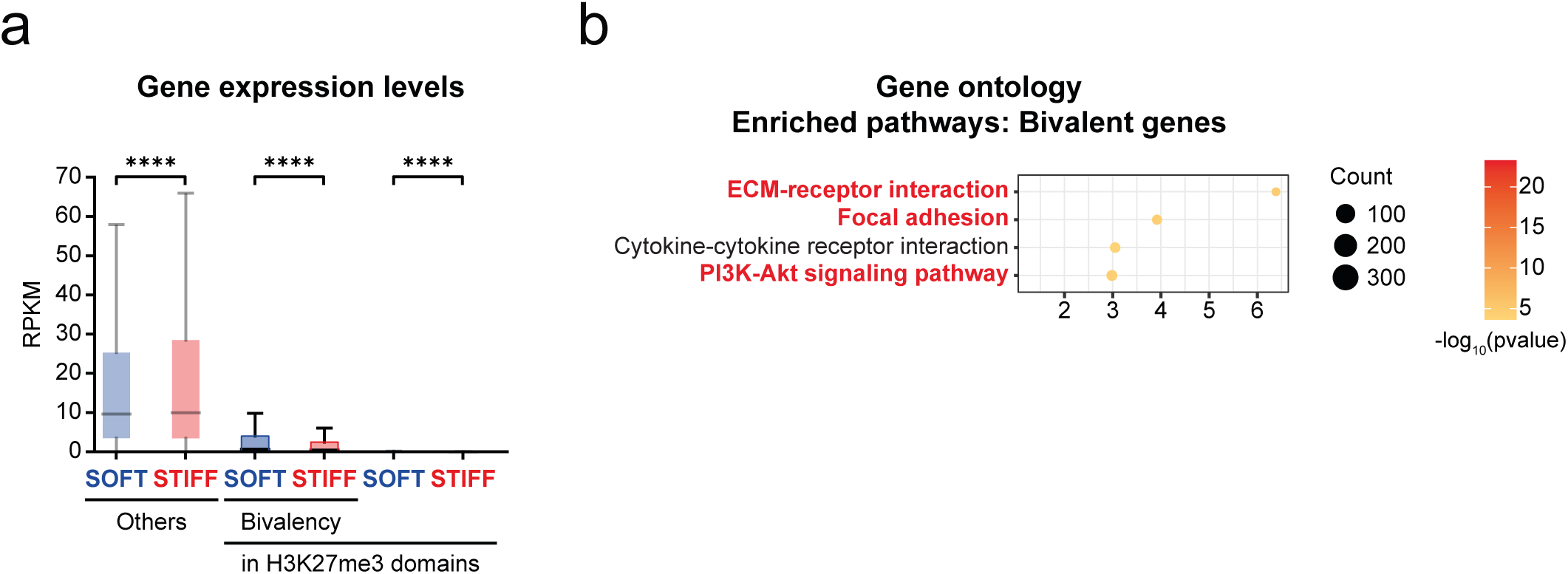
Bivalent genes tend to be more expressed under SOFT condition at genome-wide levels. a. Tukey plots of RPKM from total RNA-seq experiments for genes marked in their promoter (region -300bp before and +100bp after TSS) with H3K4me3 only (Others: SOFT n=8824 and STIFF n=8578) or with in H3K27me3 only (SOFT n=3252, STIFF n=3903). Bivalency: genes harbouring H3K4me3 and H3K27me3 at their promoters (SOFT n=606 and STIFF n=1055). Middle bar represents the median. Statistical tests are Mann-Whitney. b. Gene ontology (GO) analysis of apparent bivalent genes.

**Extended Data 4:**
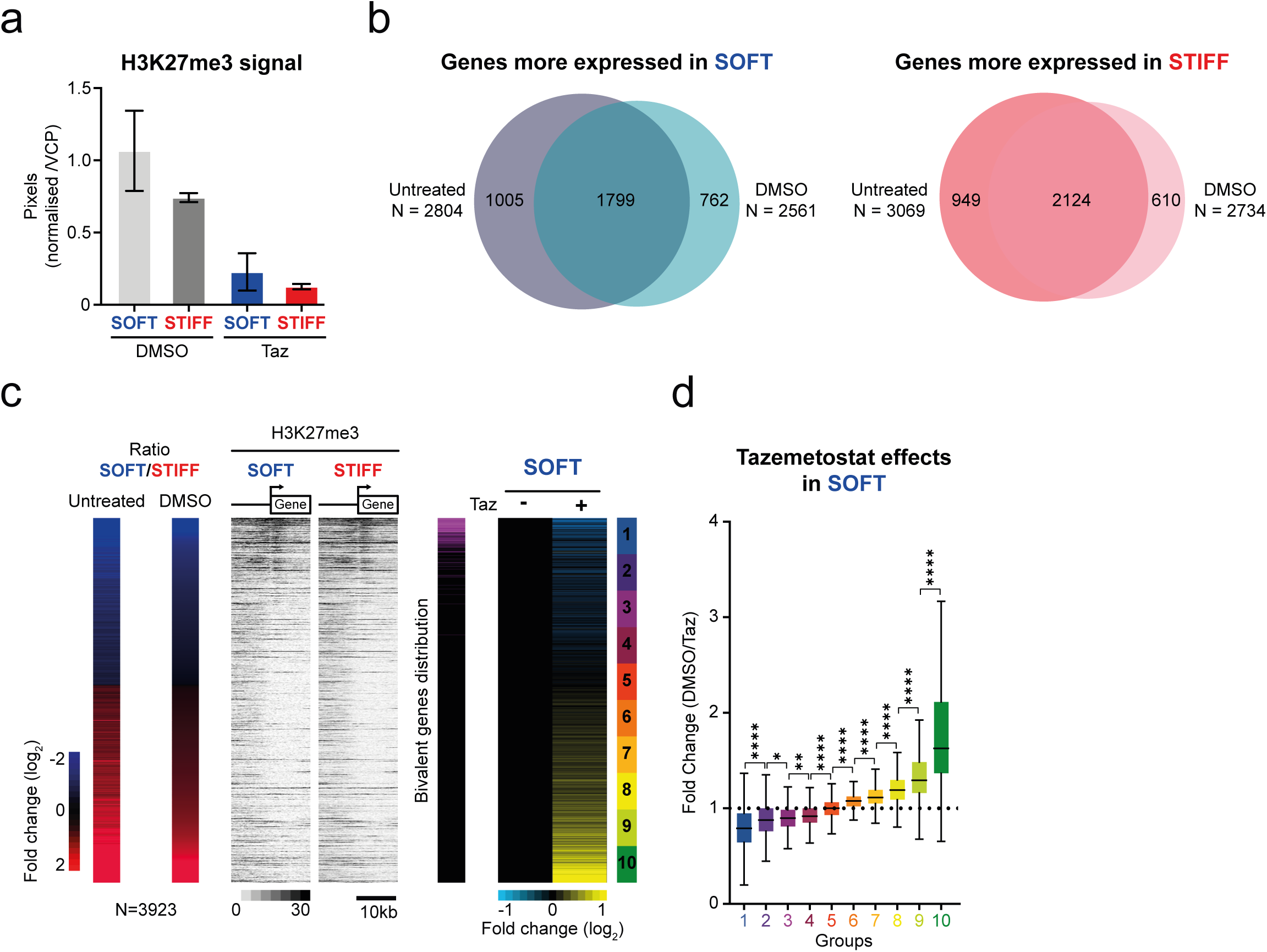
EZH2 also impact gene activation in STIFF condition. a. Quantifications H3K27me3 abundance analysed by western-blot in Figure 4c. Mean with SEM of 2 independent replicates are represented. b. Venn-diagrams comparing mechano-dependent genes identified in Untreated and DMSO condition. c. Effects of Tazemetostat treatments on gene expression of mechano-responsive genes. Only genes with common effects between untreated and DMSO control conditions were considered, to avoid indirect DMSO effects. Genes are ranked based on their total RNA fold changes between SOFT and STIFF hydrogels in the DMSO condition. Fold changes in total RNA and H3K27me3 signals in untreated condition are also displayed. Pink: distribution of apparent bivalent genes identified in **a** and **b**. Right: Tazemetostat effect on gene expression in SOFT condition in Tazemetostat treated cells (+) compared to DMSO (-). d. Tukey plots comparing fold-changes of each of the 10 groups of genes determined in **c**. Statistical tests are Mann-Whitney.

**Extended Data 5:**
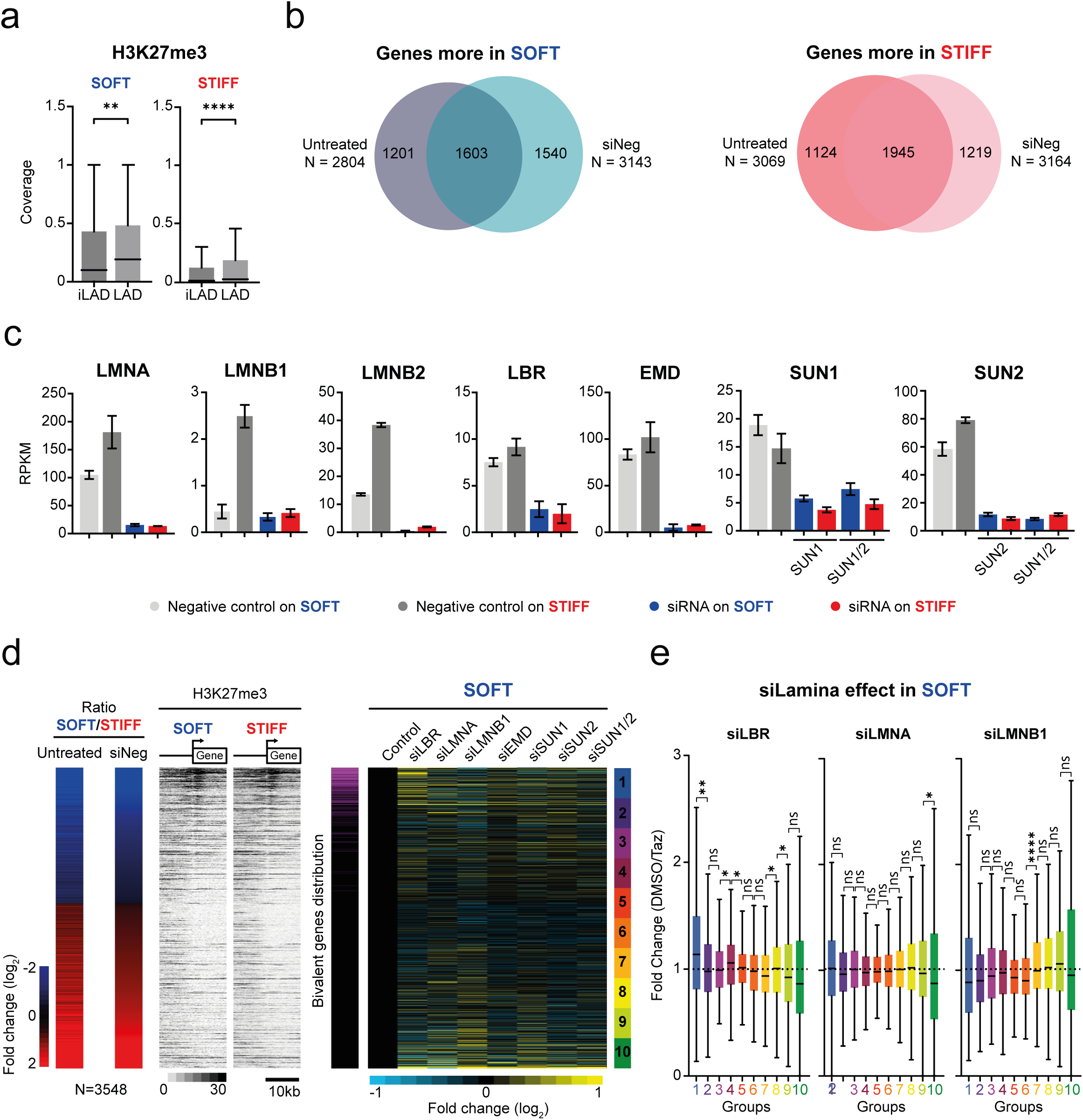
Impairing the nuclear lamina compoents has little effects on gene expression in cells grown on SOFT condition. a. Base pair coverage of H3K27me3 on interLAD (iLAD) or (LAD) depending on stiffness conditions. b. Venn-diagrams comparing differential genes identified in Untreated and siNeg conditions. c. Effects of the siRNA on their target genes. Barplots representing mean with SD of RPKM from total RNA-seq experiments of the siRNA target genes. siNeg (control) and specific siRNA effects are shown on their target genes in each stiffness condition. Each bar represents 3 independent replicates. d. Effects of siRNA treatments on gene expression of mechano-responsive genes on SOFT hydrogels. Only genes with common effects between untreated and siNEG control conditions were considered, to avoid indirect effects of transfection reagents. Genes are ranked based on their total-RNA fold change between SOFT and STIFF in the siNEG condition. Fold changes in total RNA and H3K27me3 signals in untreated condition are also displayed. Pink: distribution of apparent bivalent genes identified in **a** and **b**. Right: siRNA targeting NL or NL-associated proteins effects on gene expression in SOFT condition (+) compared to siNeg (-). e. Tukey plots comparing fold changes of each of the 10 groups of genes determined in d for siLBR, siLMNA and siLMNB1. Statistical tests are Mann-Whitney.

